# p53 amyloid pathology with cancer grades and p53 mutations

**DOI:** 10.1101/2023.07.14.547625

**Authors:** Shinjinee Sengupta, Namrata Singh, Ajoy Paul, Debalina Datta, Debdeep Chatterjee, Semanti Mukherjee, Laxmikant Gadhe, Jyoti Devi, M Yeshwant, Mohit Kumar Jolly, Samir K. Maji

## Abstract

p53 mutation and amyloid formation are implicated with cancer pathogenesis, but the direct demonstration of the link between p53 amyloid load and cancer progression is lacking. Using multi-disciplinary techniques and a cohort of 59 tumor tissues (53 from Indian cancer patients and six normal tissues) of oral and stomach cancer types, we showed that p53 amyloid load and cancer grades are highly correlated. Further, next-generation sequencing (NGS) data suggest that not only mutant p53 (e.g., SNVs, deletions, and insertions) but wild-type p53 also formed amyloids either in the nucleus (50%) and/or in the cytoplasm in most cancer tissues. Interestingly, in all these cancer tissues, p53 displays a loss of DNA binding and transcriptional activities, which is highly aggravated with the amyloid load and cancer grades. The p53 amyloids also sequester higher amounts of p63/p73 isoforms in higher-grade of tumor tissues. The data suggest p53 misfolding/aggregation and subsequent amyloid formation lead to loss and gain of p53 tumorigenic function, aggravation of which might determine the cancers grades.

## Introduction

p53 plays a vital role as a tumor suppressor and carries out different functions related to DNA repair, cell cycle arrest, apoptosis, senescence and metabolism (Aubrey et al., 2018), (Lane and Crawford, 1979), (Levine, 1997), (Mello and Attardi, 2018), (Vousden and Lu, 2002). p53 functional loss was initially known to be associated with cancer-related mutations, resulting in either destabilization of the protein fold or its inability to bind to the consensus DNA sequence (Rivlin et al., 2011), (Silva et al., 2018), (Wang and Fersht, 2015a), (Joerger et al., 2006). Mutant p53 not only displays its loss of tumor suppressor functions but also demonstrates a gain of oncogenic properties (Lang et al., 2004), (Xu et al., 2011), (Liu et al., 2010). Moreover, point mutations in any of the three domains of the p53 (N-terminal, DNA-binding, and tetramerization) lead to nuclear or cytoplasmic accumulation of p53 in cancer cells as well as in cancer tissues (Rivlin et al., 2011), (Rivlin et al., 2011), (Moll et al., 1992). The mutant p53 display the gain of functions by interacting with the wild-type p53, p53 isoforms (p63, p73), and other transcription factors (Xu et al., 2011). This results in the transcription of genes involved in cell growth, resistance to apoptosis, and metabolic reprogramming (Mantovani et al., 2019), (Moll et al., 1992). However, loss and gain of function of p53 may not always be associated with p53 gene alterations as wildtype (WT) p53 misfolding and accumulation may also contribute to cancer (Moll et al., 1992). Recent data suggest that p53 aggregation and amyloid formation is also associated with the loss of p53 tumor suppressive function and the gain of oncogenic function (Ghosh et al., 2017). Moreover, our group has recently demonstrated that p53 amyloids have prion-like properties in cells and can induce cancerous transformation in normal cells (Navalkar et al., 2020a). Although previous data suggest that p53 amyloid might be associated with cancer, the extent of p53 amyloids and cancer disease severity (such as cancer grade) is still not established yet.

In the present work, using a cohort of 59 cancer patients’ tissues with different oral and stomach cancer grades, we investigated the extent of p53 amyloid formation in different cancer grades. We found increasing p53 amyloids in higher grades of cancer biopsies for all patients for both types of cancers. Although most of these p53 accumulations are associated with p53 mutations, WT p53 accumulation and amyloid formation are also observed in certain cancer tissues. Both WT and mutant p53 amyloids are localized in the nucleus and/or cytoplasmic aggregates consisting of transcriptionally inactive p53. However, with increasing grades of both cancer types, p53 amyloids sequester more of its paralogs, such as p63 and p73, suggesting widespread deactivation of tumor suppressive function with the increase in grades due to p53 amyloid formation. Overall, the study reveals that p53 amyloid formation is not only the plausible cause of cancer initiation but may also be a positive/essential factor for cancer severity and progression.

## Results

### Increased p53 amyloid load with cancer grades

Several studies have previously suggested the formation of p53 aggregates in various tumor tissues (Ghosh et al., 2017; Moll et al., 1995; Ostermeyer et al., 1996). In this study, we examined whether p53 amyloids can be implicated as a prognosis factor for the tumor grades. For this, we used a small cohort of human cancer biopsies from Indian patients with different grades for oral and stomach cancer, which are prevalent in the Indian population. Further, among various cancers, the prevalence of p53 mutations is also high in both oral and stomach cancers (Olivier et al., 2010). We first confirmed the cancerous status of these tissues using the H&E staining (**Fig S1**). The data showed differential haematoxylin and eosin staining as well as neoplastic cells with intense nuclear staining, indicating the presence of hyperproliferative cells. To examine the p53 status and its accumulation into amyloids in cancer tissues, the double immunofluorescence colocalization study was performed using amyloid-specific antibody OC (Kayed et al., 2007) and p53-specific antibody (DO-1, Santacruz Biotechnology, Dallas, TX, USA). The data suggests the high colocalization of p53 with amyloid (OC signal) in all of the cancer tissues of oral (**Fig. 1,2**) and stomach cancers (**Fig. 3**). However, the corresponding normal tissues of the oral and stomach origin (**Fig. S2**) either showed lower levels of p53 accumulation and/or negligible p53 colocalization with OC antibody. In 59 cancer biopsies (53 tumor tissues and six normal tissues) (**Table S1**), we found that >90% of both oral and stomach tumor tissues were positive for p53 in the amyloid state. Interestingly, when analysed grade-wise manner, the p53 amyloid content (OC antibody staining) significantly increased with the progression of the tumor grades for both oral and stomach cancers (**Fig. 4A**). Similar observations were also seen when fluorescence colocalization studies were performed with p53 antibody and amyloid-specific dye ThioS staining with selected cancer tissues (**Fig. S2B-D**). Previous reports suggested p53 oligomer formation in various tumor tissues using amyloid oligomer-specific antibody A11 (Kayed et al., 2003). When we performed the double immunofluorescence study using amyloid oligomer-specific A11 antibody (red) and p53-specific antibody (green), the lower-grade cancer tissue of both oral and stomach origin showed a high degree of p53 colocalization with oligomers-specific antibody. This suggests that p53 oligomers might be formed at the initial stage of cancer, which subsequently went down in the higher tissue grades (**Fig. S3A**).

**Figure 1.**
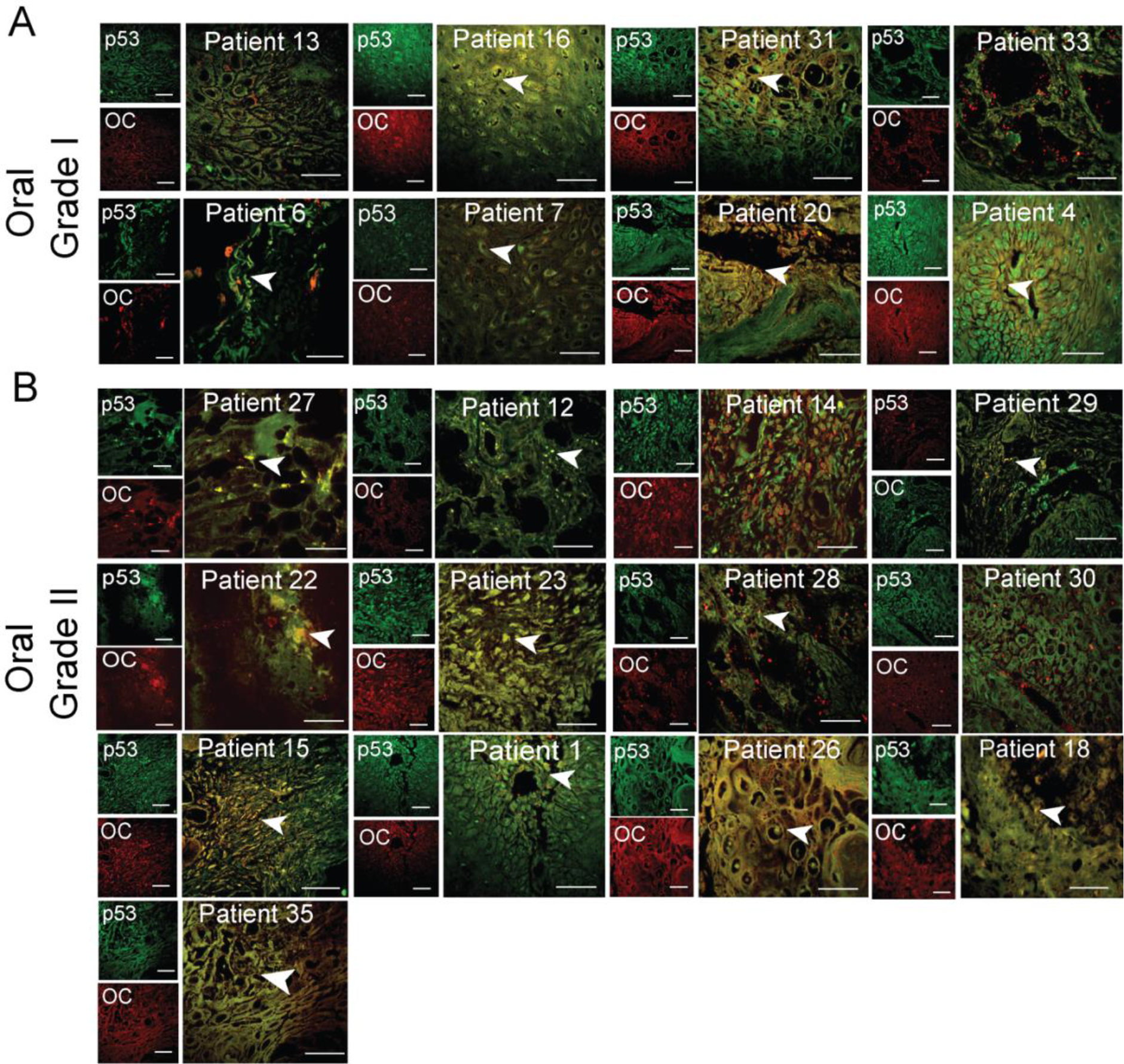
p53 status in grade I and II of oral cancer tissues. Immunohistochemistry using anti-p53 antibody, DO-1 and anti-amyloid, OC antibody showing p53 colocalization with amyloids. The yellow areas denote the colocalization of p53 and OC signals. Representative images of Oral Grade I (A) and Oral Grade II (B) were shown. The colocalized areas are shown with an white arrow. The patient number is denoted on all the images. Scale bar 50 μm. Image representative of n=2 experiments.

**Figure 2.**
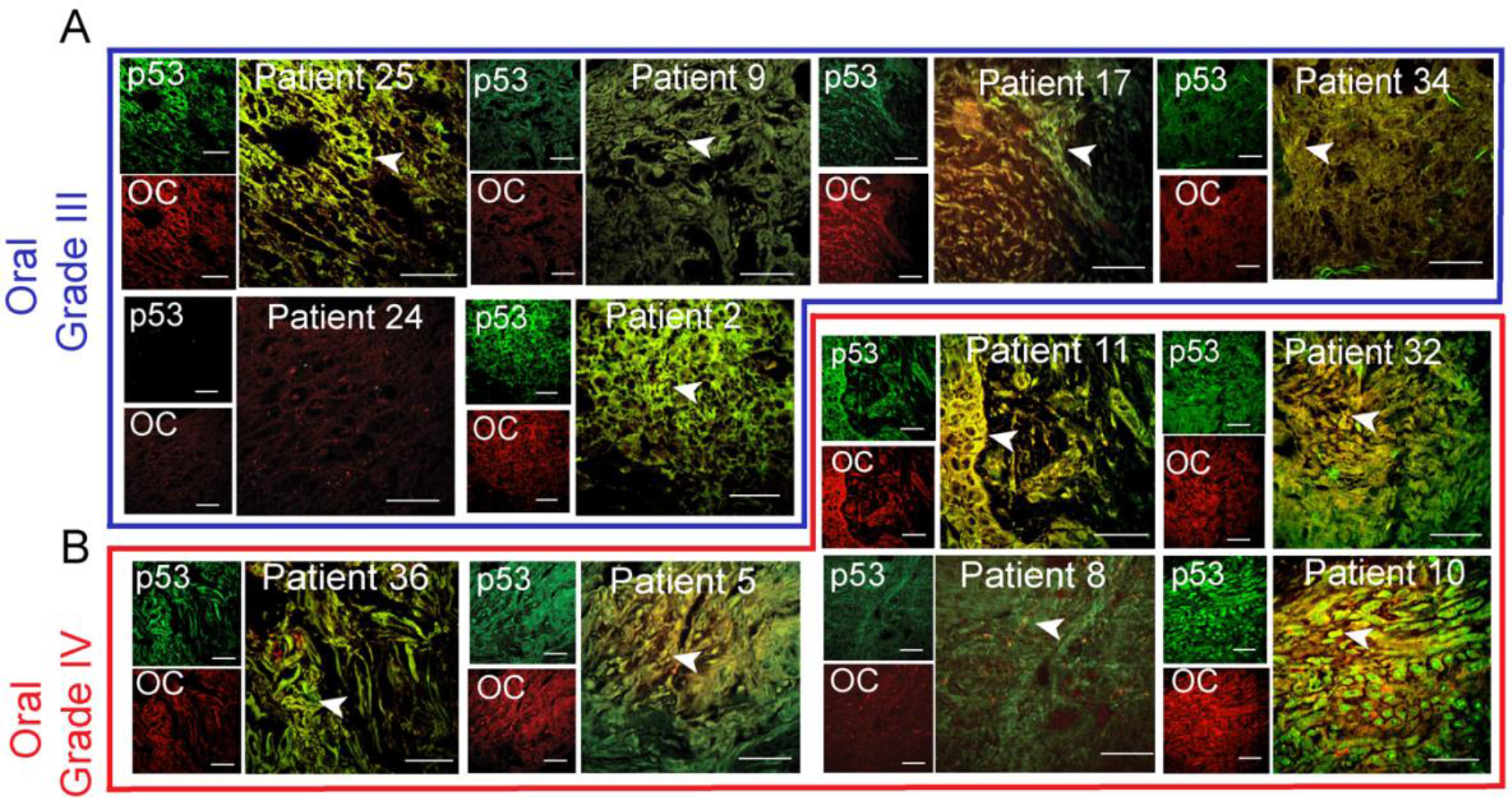
p53 status in grade III and IV oral cancer tissues. Immunohistochemistry using anti-p53 antibody, DO-1 and anti-amyloid, OC antibody showing p53 colocalization with amyloids. The yellow areas denote the colocalization of p53 and OC signals and marked with a white arrow. Representative images of Oral Grade III (A) and Oral Grade IV (B) were shown. The different oral cancer grades are highlighted in different colour boxes or lines, Oral Grade III (Blue Box) and Oral Grade IV (Red line). The patient number is denoted on all the images. Scale bar 50 μm. Image representative of n=2 experiments.

**Figure 3.**
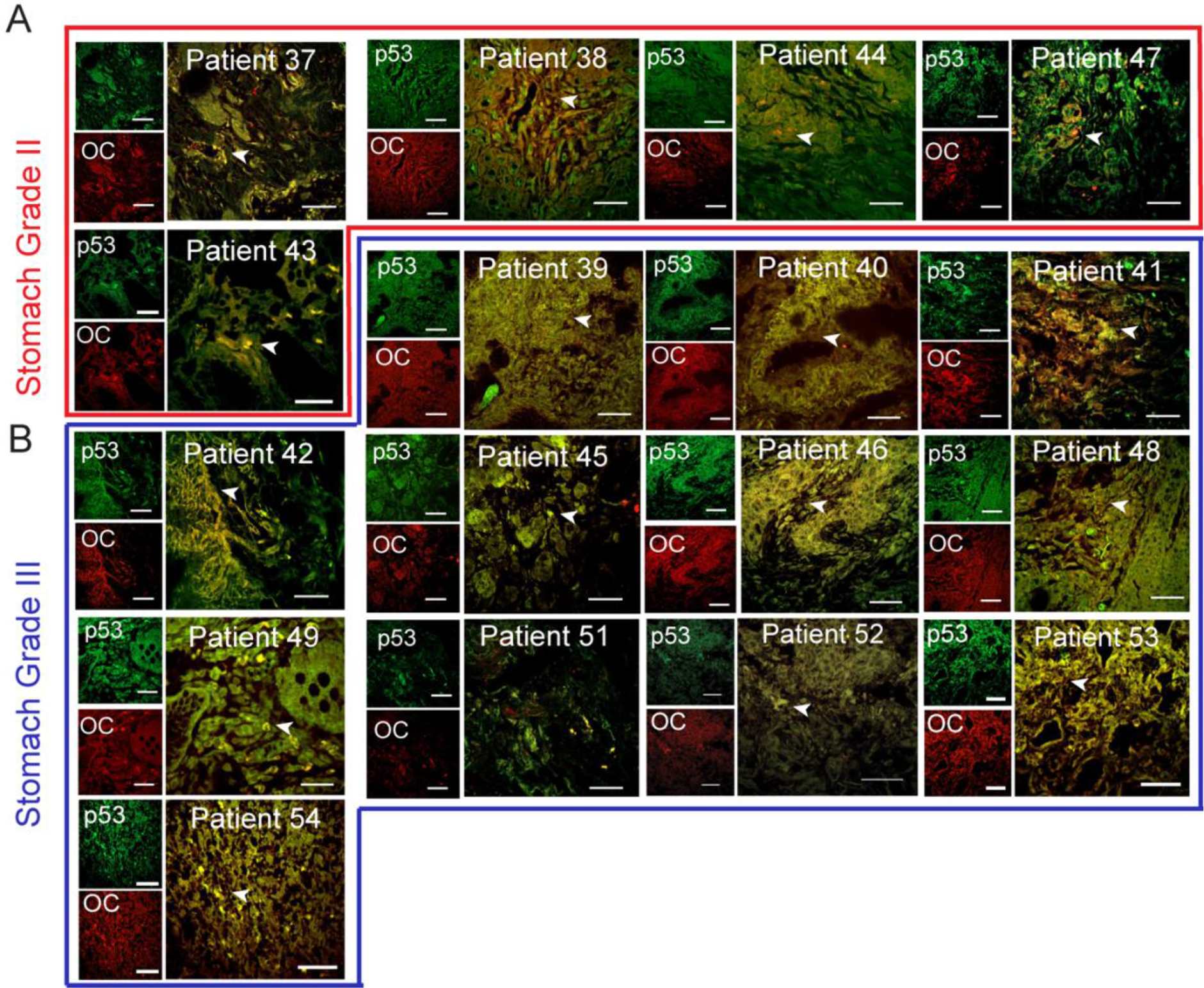
p53 status in stomach cancer tissues. Immunohistochemistry using anti-p53 antibody, DO-1 and anti-amyloid, OC antibody showing p53 colocalization with amyloids. The yellow areas denote the colocalization of p53 and OC signals and shown with a white arrow. Representative images of Stomach Grade II (A) and Stomach Grade III (B) were shown. The different stomach cancer grades are highlighted in different colour boxes, Stomach Grade II (Red Box) and Stomach Grade III (Blue Box). The patient number is denoted on the images. Scale bar 50 μm. Image representative of n=2 experiments.

**Figure 4.**
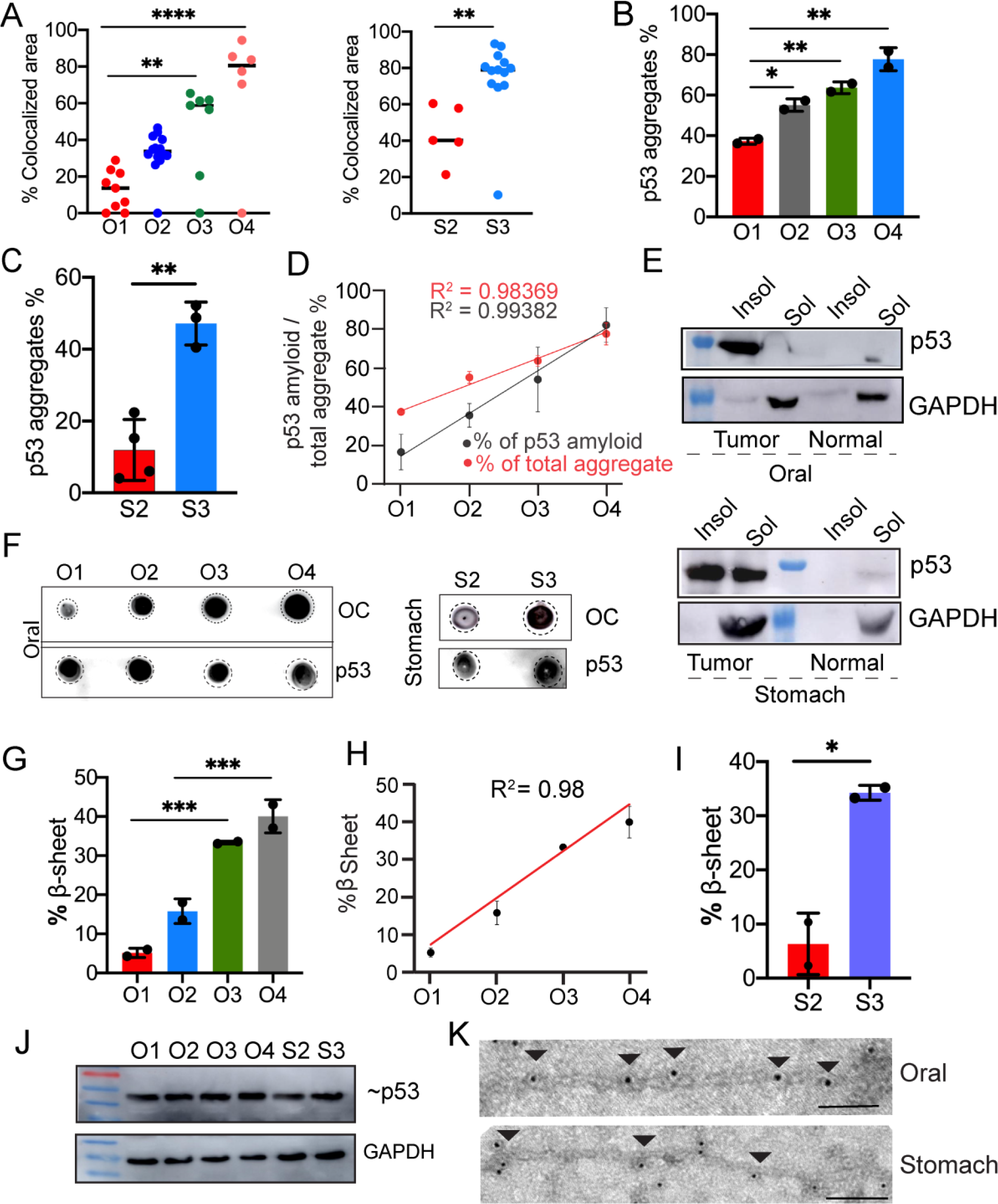
p53 amyloid characterization in oral and stomach cancer tissues. A. The colocalized areas of p53 and amyloid specific antibody OC were quantified using Image J as percent colocalized area for oral (Left Panel) and stomach (Right Panel), showing increased colocalization with higher cancer grades. The values were plotted as mean ± s.e.m., n=3 independent experiments. The statistical significance was calculated using one-way ANOVA followed by Tukey’s multiple Comparison test. B, C. Total p53 aggregate quantification using Seprion Ligand binding assay using anti-p53 antibody showing the percentage of p53 aggregation for oral cancer tissues. The values are plotted as mean ± s.e.m., n=2 and n=3 individual data sets respectively. The statistical significance was calculated using one-way ANOVA followed by Tukey’s multiple Comparison test. D. The correlation plot showing a strong positive correlation between the total p53 aggregation and p53 amyloid load in all the oral cancer grades. E. p53 expression using western blot in the soluble and insoluble fraction from oral (top panel) and stomach (lower panel) tissues showing higher content of p53 amyloid in the insoluble fraction than that of soluble fraction. GAPDH was used as a loading control. Image representative of n=2 experiments. F. Dot blot showing amyloid content and p53 expression in different grades of oral left panel) and stomach (right panel) cancer tissues. Image representative of n=2 experiments. G. H and I. FTIR image analysis of oral tumor tissues with different grades, which was quantified for the Amide I band in 1620 cm^-^ to 1640 cm^-1^ region showing an increased amount of β-sheet for a higher grade of cancer tissues. Percentage of β-sheet structure quantified from FTIR imaging of stomach cancer tissues showing the higher β-sheet structure in the higher grade of stomach cancer tissues. The values were plotted as mean ± s.e.m., n=2 independent experiments. J. p53 expression in tumor grades of both oral and stomach origin. The lower panel shows the expression of GAPDH for loading control. Image representative of n=2 experiments. K. Immunoelectron microscopy showing 10 nm gold particle decorations on the fibrils due to the presence of p53 in fibrils isolated from oral and stomach cancer tissues. Scale bars, 200 nm. Images representative of n=3 independent experiments.

Further, to quantify the total p53 aggregation in various grades of cancer tissues, we used an enzyme-linked-immunosorbent-assay (ELISA) based on a polyionic, high-molecular-weight ligand that specifically binds to aggregated proteins (Maritschnegg et al., 2018). Consistent with the double immunofluorescence data (**Fig. 1,2 and 3**), the levels of p53 aggregates were increased many folds in the higher-grades of oral and stomach cancer tissues, compared to the corresponding lower grades (**Fig. 4B**, **Fig. 4C**), exhibiting a positive correlation in oral cancer tissues (R^2^= 0.98) (**Fig. 4D**). We found high correlation of the total p53 aggregates (ELISA) and p53 amyloid (OC staining) (**Fig. 4D**). Interestingly, we further observed that in lower grades, p53 amyloid pool is much lesser than total p53 aggregate, however with higher cancer grade, p53 amyloid predominates exhibiting positive correlation (R^2^= 0.98 for total p53 aggregates and R^2^= 0.99 for total p53 amyloid) (**Fig. 4D**). The higher amount of p53 amyloids in higher cancer grades was also consistent with western blot analysis of soluble versus insoluble p53 fractions (**Fig. 4E, Fig S3B,C**) and dot-blot analysis (**Fig. 4F and Fig. S3D,E**) from lysate isolated from various grades of cancer tissue extracts. Important to note that, In normal tissues, only faint band of p53 in soluble fraction was observed. This is well-known fact that p53 is not detectable in normal tissue and cells as it readily degrades and negatively regulated due to MDM2 (Marine and Lozano, 2010; Francoz et al., 2006).

Overall, the data suggest that increased extent of p53 amyloid formation in a higher grade of oral and stomach cancers. This high extent of p53 amyloid formation at the higher stage of cancer could be due to widespread misfolding and amyloid amplification of p53. We further intended to examine whether higher p53 amyloid loads can be correlated with other cancer grades using bioinformatic analysis. For this, we hypothesized that if p53 amyloid load is correlated with cancer grades, it should result in a greater extent of altered gene expression patterns associated with p53 amyloids. In this context, our previous study showed uniquely altered gene expressions associated with p53 amyloids in cells in contrast to the normal cells and p53 cancer-associated mutations (Navalkar et al., 2021; Navalkar et al., 2022). We first analysed the correlation of these altered genes (associated with p53 amyloids) in Head and neck cancer tumor samples using UALCAN database. We found the enrichment of these unique genes are well correlated with cancer grades (**Fig. S4A)**. Similar observations were also obtained with other various cancers (**Fig. S4B-G**). The data, therefore, clearly showed that a higher amount of p53 amyloids are directly associated with the higher grade of cancers and supports the idea that p53 amyloids act as an oncogene for promoting cancer pathogenesis.

### Characterization of p53 amyloid in cancer tissues

To characterize the amyloid content of the various cancer tissues, we performed Fourier-transform infrared (FTIR) imaging (Miller et al., 2013). The FTIR imaging with snap-frozen tumor biopsies showed a higher amount of β-sheet content in higher-grade oral cancers compared to the corresponding lower-grade and normal tissues (**Fig. 4G, 4H, 4I, Fig. S5, and Fig. S6**). Similar observations were also obtained for stomach cancer biopsies (**Fig. 4I**). Important to note that the higher β-sheet content could be due to other protein amyloids along with p53 amyloids. However, our combined study of immunohistochemistry and label-free FTIR imaging on the identical tissue section (used adjacent sections for both the study) support that the presence of p53 amyloid might be mostly responsible for higher β-sheet-rich structure in these tissue sections (**Fig. 4 G-I; Fig. S5, and 6**). However, there is a possibility that amount of p53 expression/stabilization might be higher in higher grade of cancer compared to lower grade. To examine this, western blot analysis is done for 2 patients from each grade (I,II,III and IV) tissue for oral and 2 patients from each grade tissue for stomach (II and III). The data showed similar expression of p53 from grade I towards the higher grade cancer (**Fig 4J, Fig. S7A**). As our double immunofluorescence data showed that in the lower grade of cancer tissue, p53 amyloid oligomers are present but not in the higher grade. Therefore, amount of total p53 might be same but their state/conformation might differ among grades where p53 amyloid dominates in the higher grade of cancer tissues.

We further isolated the tissue amyloid fraction (TAF) from representative human oral cancer and stomach cancer tissues, which showed fibrillar morphology under the electron microscope (EM) (**Fig. 4K**). FTIR study of these p53 fibrils showed the presence of intense peaks at ∼1627 for oral and ∼1632 for stomach cancer in the amide I region (**Fig. S5, and Fig. S6**) suggesting the structure of the β-sheet-rich amyloid (Jackson and Mantsch, 1995) in these p53 aggregates. Indeed, the immunoelectron microscopy (immuno EM) using p53 antibody (primary) and 10 nm colloidal gold-conjugated secondary antibody confirmed p53 amyloid as gold particles were aligned along the length of isolated fibrils from oral and stomach cancer (**Fig. 4K and Fig S5C**).

### Wild type (WT) and mutant p53 are responsible for amyloid formation in tumor tissues

p53 mutations are known to be associated with 50% of human cancers (Soussi et al., 2006). Most often, p53 mutations result in the accumulation of p53 as a punctate appearance in various cancer cells and tissues (Moll et al., 1995). Further, it was shown that mutant p53 preferentially accumulates in the nucleus, whereas WT p53 sequesters in the cytoplasm of the cancer cells (Moll et al., 1995). These p53 accumulations are either due to WT p53 destabilization, which could be further induced by p53 mutations (Kim et al., 2009), (Soussi et al., 2006), (Wang and Fersht, 2015a). Since we observed grade-wise increase in p53 amyloid formation (**Fig. 4**) in cancer tissues, we examined whether there is any link between p53 amyloids and p53 mutations. For p53 mutational status, we performed Next-Generation Sequencing (NGS) for 48 cancer tissues (44 tumor and 4 normal tissues) (**Fig. 5, Table S1**), which showed Tp53 mutations in ∼93% of the tumor tissues (**Table S1**). Several SNVs, deletions, insertions, and stop-gain mutations were observed in both these tumor tissues. Although, the highest mutation type observed was single nucleotide variants (SNVs) in the p53 gene in both oral and stomach cancer tissues. Interestingly, SNVs including various hotspots mutations (R175H, E286V, R267W, R248W, R282W, R248L, and E285K), stop-gain mutations (such as R306/*, R196/*, Q317/* and R213/*), insertion and deletions were mostly detected in the DNA binding domain of TP53 gene (94-312 amino acids) (**Fig. 5A, and Table S1**) in oral cancer tissues. In the tetramerization domain (325-356), only one SNV (R337C), along with a stop-gain mutation R342/* and no mutations were detected in the transactivation domain of the p53 gene (**Fig. 5A, and Table S1**). This is consistent with previous p53 mutation data, which suggest that the frequency of p53 mutations mainly occurs in p53 DBD (Rivlin et al., 2011). We further found that the missense mutations in the DNA binding domain were highest at exon 4 followed by exon 8 (**Fig. 5B, E**). When we analyzed the extent of p53 mutations in various cancer grades, we observed an increased extent of SNV mutations in a higher grade of cancer tissues (Moll et al., 1995), (Liu et al., 2010), (Skinner et al., 2012). This suggests that p53 mutations could be one of the primary factors for the destabilization and amyloid aggregation of p53 (**Fig. 5C, D, and E**). Our results provide support for the correlation between p53 SNVs and amyloid formation in different cancer grades, with the exception of oral cancer grade III, which exhibited a minimal number of SNVs. (**Fig. 5D**). Previous data indicates that certain p53 mutations, such as R175H, R248W, and R282W, have a higher likelihood of causing misfolding and aggregation of the p53 protein. (Ghosh et al., 2017), (Palanikumar et al., 2021), (Ferretti et al., 2022). A similar mutational status was also observed in the case of stomach cancer tissues where SNVs (G245V, I232F, Y220H, A138S, and C176F), deletions, and insertions were frequently observed in the DNA binding domain. We, however, observed neither stop-gain mutations nor SNVs in other domains of the p53 gene. Further, the transition of G>C was observed to be the highest in these tissue samples (**Fig. 5F**) compared to other transitions such as G> A, C>A, T>A, and A>G. Important to note that many of these mutations that we observed are already known to be directly associated with human cancer (Mello and Attardi, 2013), (Monti et al., 2020), (Moll et al., 1995), (Bauer et al., 2020), (Manterola et al., 2018). However, for some mutations (e.g., P72R), further studies are required for their role in cancer pathogenesis (**Fig. 5G**). Although most of the cancer tissues containing the p53 amyloids possess frequent mutations in p53 gene, we, however, observed p53 amyloids with wild type protein in cancer tissues (patient number 35, 38 and 42). The data indicates that the misfolding, and aggregation followed by amyloid formation by p53, which is further promoted by cancer-associated mutations, might result in a higher amount of p53 amyloids in a higher grades of cancer tissues.

**Figure 5.**
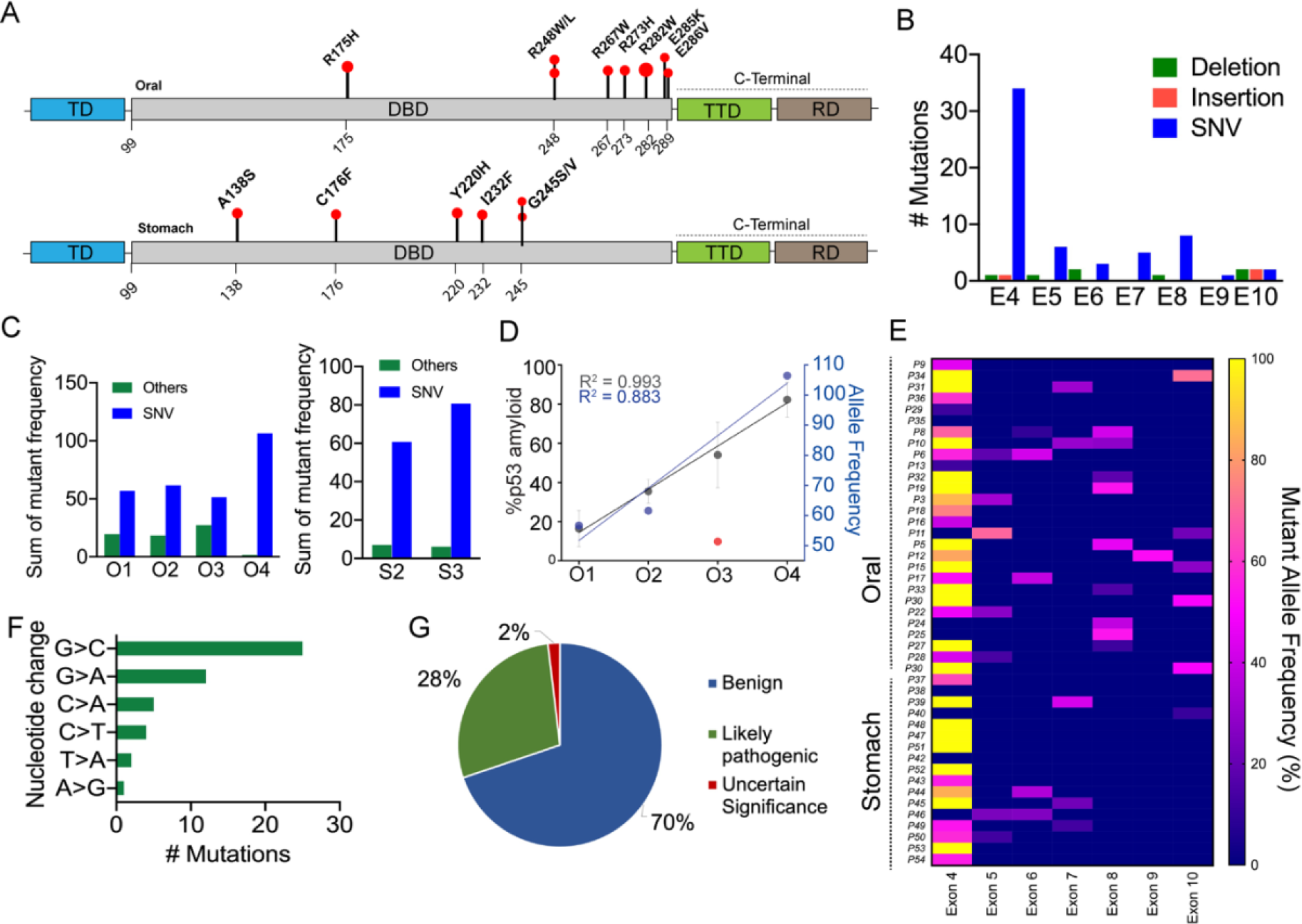
Detection of mutations in human oral and stomach biopsies by next-generation sequencing. A. Schematic of p53 domains with lollipop plot showing the single nucleotide variants observed in the oral biopsies (upper panel) and stomach biopsies (lower panel). B. Cohort samples (n=48) showing the total mutations observed in different exons of the p53 gene. C. Alternate allele frequency showing variation in mutation types for different grades of cancer tissues. D. Correlation plot showing percent p53 amyloid and alternate allelic frequency of SNVs in different oral cancer grades. E. Heat map of all the cohort samples showing the mutation frequency in different p53 exonic regions. F. Mutation frequency among different patients in the cohort showing nucleotide substitution. G. Pie chart showing the clinical significance of the NGS detected mutations as obtained from NCBI database search.

### Cytoplasmic versus nuclear sequestration of misfolded p53 in tumor tissues

On the onset of stress (e.g., DNA damage), phosphorylated p53 enters into the nucleus to carry out the transcriptional function by binding to its cognate DNA sequence (Sammons et al., 2020). However, p53 can be excluded from the nucleus (Lu et al., 2000) and shuttle between cytoplasm and nucleus where the nuclear localization signal (NLS) and nuclear export signal (NES) play an essential role for this nucleocytoplasmic transportation (Liang and Clarke, 2001). Previous reports suggest that mutant p53 accumulates preferentially to the nucleus, whereas wild-type p53 protein is sequestered and stabilized into the cytoplasm, thus rendering it non-functional (Moll et al., 1992).

Since we examined p53 amyloids in a relatively large number of cancer tissues with different grades and established the p53 mutational status, we examined the correlation between the mutational status of p53 and the location of p53 amyloids. For this, we used DAB (3, 3’-diaminobenzidine) staining for all tumor biopsies using a Pab240 monoclonal antibody (Santacruz), which recognizes misfolded p53 under non-denaturing conditions (**Fig 6, Fig. 7A,B,C**). However, to examine whether DAB staining indeed recapitulates the p53 amyloids, we parallelly examined co-immunofluorescence with Pab240 and amyloid-specific OC antibody or amyloid-specific dye ThioS (**Fig. 7A,B,C *Right panel*; Fig. S7C-E**) with selected tissues of different grades. The colocalization experiments with Pab240 antibody and OC antibody or ThioS dye (**Fig. S7C-E**) revealed colocalization of misfolded p53 with OC antibody in all cancer tissues (**Fig. 7A, B, and C**) but not in the corresponding normal tissues (**Fig S7B**). Thus, DAB staining along with p53 mutational status will allow us to understand the effect of p53 mutations on the localization of amyloid p53 in cancer tissues. The localization of p53 in selected biopsies (nuclear localization of p53 by immunofluorescence) was further confirmed by dot blot (**Fig. 7E**) analysis of nuclear and cytoplasmic extracts of these patients’ biopsies (P1 and P39). The DAB staining of cancer tissues showed three distinct nucleo-cytoplasmic staining patterns (**Fig. 7A,B,C**). Eighteen oral patients biopsies out of 36 patients (∼50 %) and seven stomach biopsies out of 17 patients (∼ 41 %) displayed only nuclear staining suggesting the accumulation of high levels of misfolded p53 protein in the nucleus (**Fig. 7**). Only cytoplasmic accumulation of p53 was observed in four oral biopsies out of 36 patients (∼ 11 %) and five stomach biopsies out of 15 patients (∼33 %). In all other patients’ biopsies (twelve oral (∼33 %) and five stomach biopsies (∼ 29%)), we observed p53 to localize in the nuclear as well as in the cytoplasm. When p53 localization is compared with mutational status, we observed that hot spot mutations either accumulate in the nucleus (such as R337C, R267W, and R248W in oral cancer) or in both nucleus and cytoplasm (R175H and G245V). Next, we analysed the extent of p53 localization in the nucleus versus cytoplasm in all the cancer tissues of pab240 antibody staining using Image J (**Fig. 7D**). Our data reveal that there is no correlation between p53 mutation and their sequestration either in the nucleus or in the cytoplasm as both WT and mutant p53 are seen to be sequestered in both nucleus and/or cytoplasm (**Fig. 7D**).

**Figure 6.**
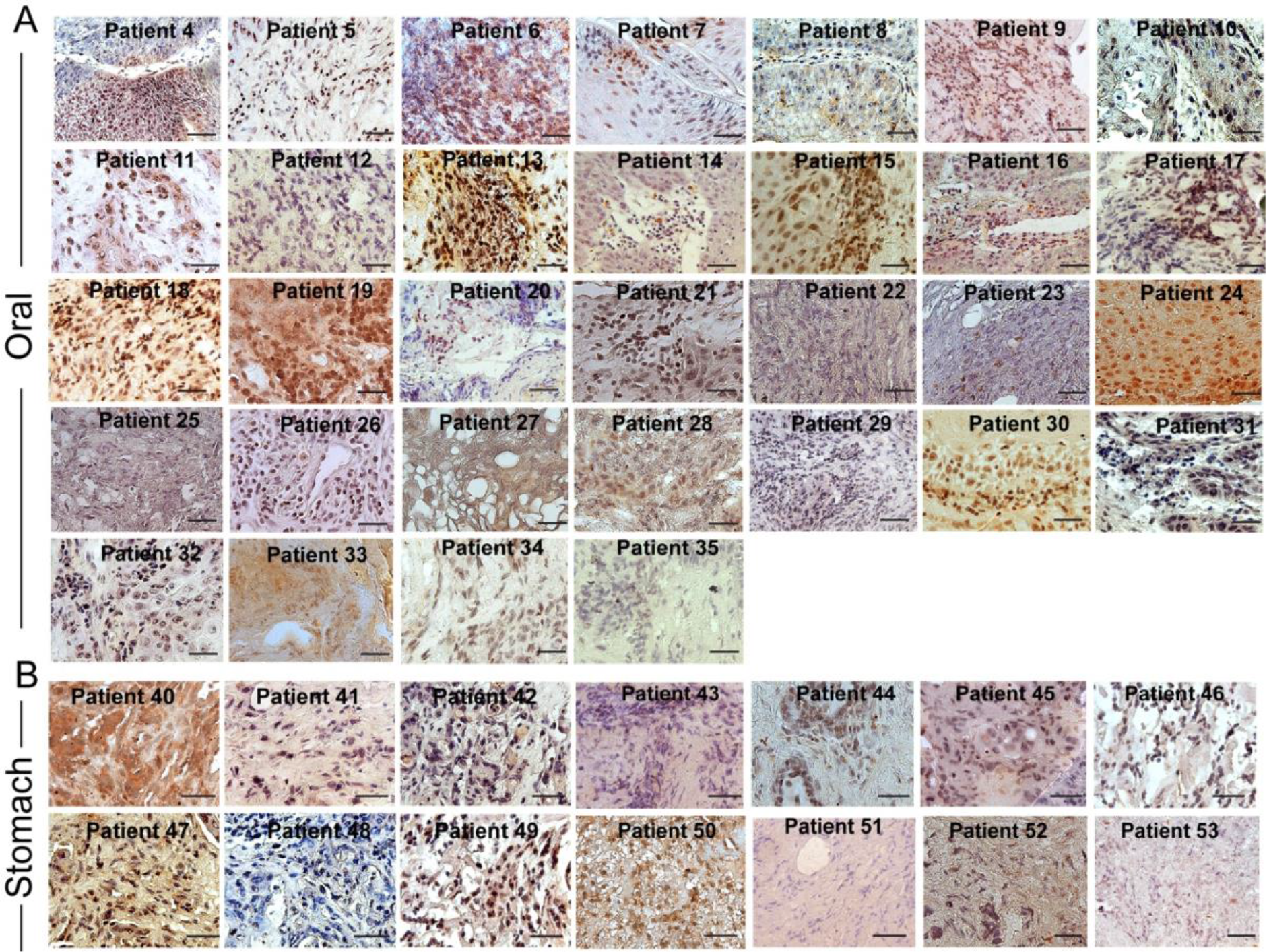
Nuclear versus cytoplasmic inclusion of p53 in oral and stomach cancer tissue biopsies. Immunohistochemical study showing misfolded p53 inclusion in (A) oral and (B) stomach cancer tissues using Pab240 antibody (Santa Cruz Biotechnology, Dallas, TX, USA) and subsequently developed by 3’-Diaminobenzidine (DAB) substrate (dark brown to light brown due to binding affinity). The tissues were counterstained with Harris Haematoxylin (Blue/purple color). The data showing nuclear or cytoplasmic expression of p53 in various cancer tissues. Scale bars are 50 μm. Images representative of n=3 independent experiments.

**Figure 7.**
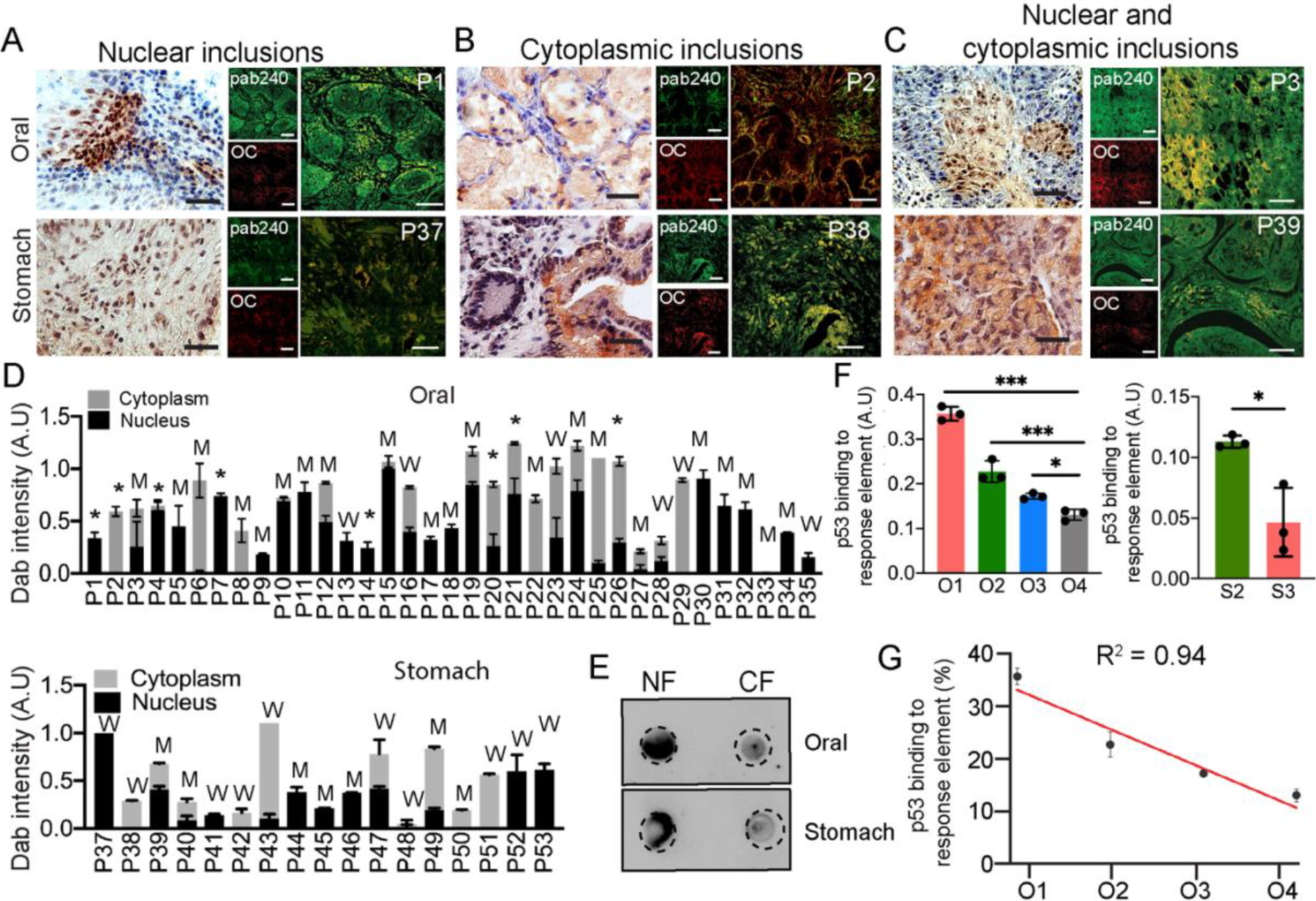
Nuclear and cytoplasmic localization of p53. Immunohistochemical staining (DAB) for misfolded p53 using Pab240 antibody showing the representative image of nuclear inclusions (A), cytoplasmic (B), and nucleo-cytoplasmic inclusion (C) of cancer tissues. Corresponding p53 amyloids were shown using colocalization by Pab240 antibody and amyloid-specific OC antibody. Representative images of both the oral (upper panel) and the stomach (lower panel) are shown. Images in A,B and C are representative of n=3 independent experiments. D. Dab intensity quantification showing the relative extent of misfolded p53 localization in nuclear, cytoplasmic, and nucleo-cytoplasmic fractions as measured using Image J for individual oral (upper) and stomach (lower) cancer tissues. The data was plotted as a stacked bar plot with the average dab intensity of one patient. The W denotes wildtype p53 in cancer tissues, M denotes mutated p53 (SNVs, deletions, or insertions), and * denotes those tissue samples not analysed by next-generation sequencing. Results are given as mean±s.e.m (n=3). E. Dot blot using Pab240 showing higher load/inclusions of misfolded p53 in nuclear than in cytoplasm in nuclear-localized patient tissue. F. p53 DNA binding ability using tissue lysate of different grades of oral tissues (Left panel) and stomach tissues (right panel) showing significant loss of p53 binding to DNA in higher grades of cancer. Results in C,D are given as mean ± s.e.m (n=3). G. The correlation plot of p53 activity with oral cancer grades shows a decrease in p53 DNA binding ability with an increase in cancer grades.

To further analyse whether these aggregated and misfolded p53 have the DNA binding ability, we performed ELISA assay using tissue lysates from 9 oral (Left Panel) and 3 stomach cancer patients’ biopsies (Right Panel) (**Fig. 7F**). The 3 tumor biopsies were chosen from each cancer grade. In all these cases and irrespective of p53 localization, the sequestered p53 was observed to be transcriptionally inactive due to their inability to binding to DNA. We also observed the much lesser DNA binding capacity (higher p53 inactivation) in higher grades of cancer (**Fig. 7F,G**). Important to note that one of the stomach cancer tissue samples harbouring wildtype p53 amyloid also displayed no p53 DNA binding capacity, suggesting their transcriptional inactivation. Therefore, the data suggest that irrespective of localization/mutation, the higher amount of p53 accumulation into amyloids resulted in p53’s inability to bind to cognate DNA sequence in the higher grade of cancers.

### p53 loss of function and higher sequestration of p63/p73 with p53 amyloids in higher grade of cancer

p53 amyloid formation has been shown to exhibit loss of tumor suppressive function as well as gain of oncogenic properties (Ghosh et al., 2017; Navalkar et al., 2021; Navalkar et al., 2022), (Navalkar et al., 2020b), (Sengupta et al., 2022). As a transcription factor, p53 recognizes its target genes by binding to a consensus response element located at the gene promoter (Rivlin et al., 2011). To examine the loss of function due to the gradual increase of p53 amyloids in higher grades of tumor tissues, we analysed 6 oral and 6 stomach cancer patients’ tumor biopsies (3 patients each from cancer grade I and grade III of oral cancer and 3 patients each from cancer grade II and III of stomach cancer) by chromatin immunoprecipitation (CHIP) assay using anti-p53-DO1 antibody (primer sequence listed in **Table S2**). With increased p53 accumulation (amyloids) from grade I to III, we observed a reduction in the extent of p53 bound with response elements (responsible for apoptosis or cell cycle arrest such as p21, PIG, Gadd45) (**Fig. 8A,B**). In contrast, lower-grade tumors containing a low amount of p53 accumulation (amyloids) showed a high amount of p53 bound with response elements (**Fig. 8A,B**). The observation is also further confirmed by quantitative qPCR analysis using the p21 gene (**Fig. 8B**). The data suggests that more p53 accumulation as amyloids results in less amount of p53 bound to its response element in higher cancer grades. We further hypothesized that widespread p53 inactivation as a tumour suppressor and gain of oncogenic functions at a higher grade of cancers might happen not only due to the gradual increase of p53 amyloids but also sequestration of other tumor suppressor proteins (e.g., p63 and p73) by p53 amyloids. In this context, it has been shown that p53 aggregates sequestered p53 paralogs such as p63/p73 in cells (Xu et al., 2011). Further, p63 and p73 are known to be rarely mutated in tumors; however, their tumor suppressor functions are frequently inhibited by mutant p53 (Inoue and Fry, 2014). To examine the sequestration of p63/p73 in p53 amyloids in these cancer tissues, co-immunofluorescence experiments were performed using p53 with p73 or p63 antibodies. We analysed 12 oral and 6 stomach cancer patients (3 patients from each cancer grade). p53 was observed to colocalize with p73 (**Fig. 8C**) and p63 (**Fig. 8D**) in all the cancer grades for both oral and stomach cancer biopsies. The image J analysis suggests the percentage of colocalization was significantly greater in the higher cancer grades for both oral and stomach (**Fig. 8 E-H**). In oral cancer, the colocalization of p53 and p73 was ∼30% in grade I, which increased up to 70% in grade IV suggesting p53/p73 colocalization is highly correlated with oral cancer grade (**Fig. 8F**). Similar observations were also seen between stomach grade II (20% colocalization) and grade III (60% colocalization). Similar to p53/p73 colocalization, we also observed a higher degree of colocalization of p53/p63 as an increase in cancer grades for both oral and stomach cancer tissue. The colocalization of p53/p63 was ∼20-25% in oral and stomach Grade I tissues, which increased to ∼ 60% for both oral grade IV and stomach grade III tissues (**Fig. 8 G-H**).

**Figure 8.**
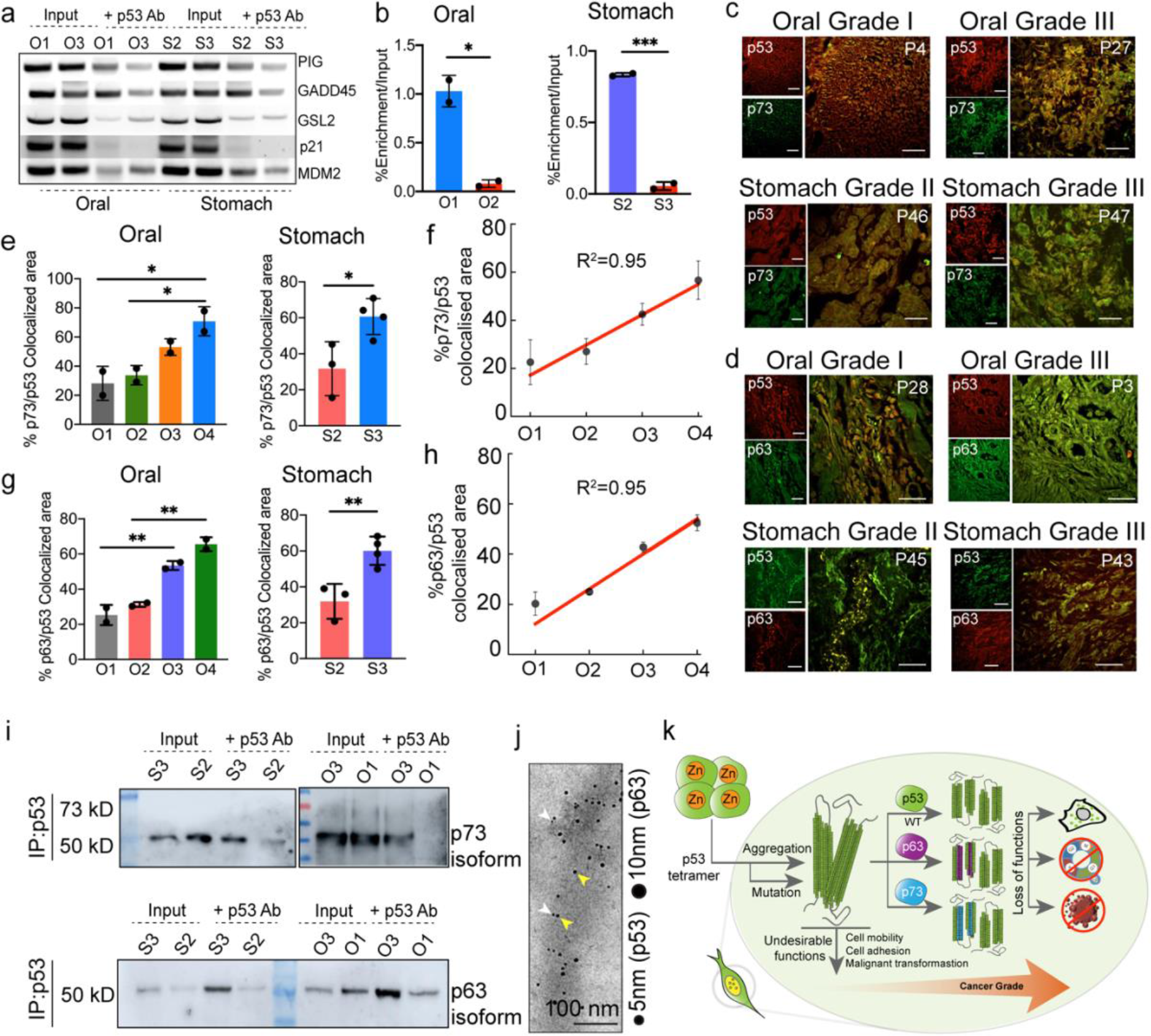
Loss and gain of function by p53 due to p53 amyloids in cancer tissues. A. Chromatin immunoprecipitation (ChIP) assay showing increased loss of p53 binding to its response element in higher grades when compared with lower grades. p53, showing its inability to bind to the response elements of PIG, GADD45, GSL2, P21 & MDM2. Input represents the whole-cell extract. Image representative of n=2 experiments. Statistical values were calculated using an unpaired test. B. Quantitative real-time PCR verifying the functional loss of p53 from oral (left panel) and stomach (right panel) tissues showing its inability to bind to p21 promoter in a higher grade of cancer. The values were plotted as mean ± s.e.m., n=2 independent experiments. C. Double immunohistochemistry (using anti-p53 antibody, DO-1 and anti-p73 antibody) showing colocalization of p53 and p73 signals in oral cancers of different grades (O1, O3, upper panel) and stomach cancer (S2, S3, lower panel). The colocalization is higher in higher cancer-grade tissue. Images shown are from n=3 independent experiments. D. Double immunohistochemistry showing colocalization of p53 and p63 signals in oral cancers (O1, O3, upper panel) and stomach cancer (S2, S3, lower panel). Image representative of n=3 experiments. E. The quantification of p53/p73 colocalization using image J showing a significant increase in the extent of co-aggregation with increased cancer grades for both oral (left) and stomach (right panel). The values were plotted as mean ± s.e.m., n=2 independent experiments. F. The plot showing a high correlation between the percentage of p53/p73 colocalization with oral cancer grades. G. The quantification of p53/p63 colocalization using image J showing a significant increase in the co-aggregation with an increase in cancer grades for both oral (left) and stomach (right panel). The values were plotted as mean ± s.e.m., n=2 independent experiments. H. The correlation plot showing a high positive correlation between the percentage of p53/p63 colocalization with oral cancer grades. I. Western blot analysis of immunoprecipitated p53 from the stomach and oral cancer tissues using p73 and p63 antibodies showing the presence of p73/p63 isoforms in immunoprecipitated p53. The amount of p73/p63 isoforms was higher in the higher cancer grade than in the corresponding lower grade. J. Co-immunoelectron microscopy showing 5 nm (gold labelled secondary antibody against primary antibody of p53) and 10 nm gold (gold labelled secondary antibody against primary antibody of p63) particle decorations on the fibrils due to the presence of p53 and p63 co-aggregation in fibrils isolated from grade III oral cancer tissues. Scale bars, 200 nm. Image representative of n=2 experiments. K. The schematic showing p53 amyloid load is corelated with the increase in cancer grades. p53 amyloid can sequester other family members, p63 and p73 which are higher in higher cancer grades. p53 amyloid results in loss of function and gain of oncogenic properties of p53.

Next, to directly confirm the p53/p73 and p53/p63 co-aggregation, p53 was immunoprecipitated using p53 DO-1 antibody from oral biopsies of grade I (P4), III (P27)) and stomach biopsies of Grade II (P46), III (P47) followed by western blot analysis with p73 and p63 antibodies. The Western blot signal confirms the presence of p73 and p63 isoforms along with immunoprecipitated p53 in higher grade (grade III) for both oral and stomach tissues (**Fig. 8I, Fig. S8A-E**). Based on the molecular weight of these proteins, we conclude that these are isoforms of p63 and p73 that could possibly be co-aggregating with the p53 fibrils in both the cancer tissue types as a gain of function property displayed by the p53 amyloids (**Fig. 8I, Fig. 8K**). However, in some of the patient samples, we also observed faint expression of full length p73 along with immunoprecipitated p53 in both oral and stomach samples (**Fig. S8C**). Further to examine the co-immunoprecipitation of p53/p63 and p53/p73, we also immunoprecipitated using antibody of p63 and p73 and then performed western blot with anti-p53 antibody (DO1). We indeed found similar observation that along with both p63 and p73, we found the presence of p53 (**Fig. S8D**). This suggest that p63 and p73 co-aggregates with p53. We further confirm this using double immunoelectron microscopy of an oral cancer tissue (grade III) with anti-p53 antibody (DOI) and anti-p63 antibody. We found both species are there in a single fibrils (**Fig 8J, Fig. S8E**). Moreover, important to note that due to the higher colocalization, at this moment, we couldn’t examine whether p63 or p73 are also in amyloid state using OC or ThioS staining in cancer tissues. Future study is required to determine whether p63/p73 alone can form amyloid independent of p53, which can be associated with cancers.

## Discussion

Amyloid formation is generally associated with neurodegenerative diseases, such as Parkinson’s, Alzheimer’s, and prion disease (Chiti and Dobson, 2017), (Dobson, 2001). Due to amyloid formation, there is a loss of particular protein function, and the gain of toxic function, which leads to cell death and neurodegeneration (Eisenberg and Jucker, 2012). In contrast to disease-associated amyloid, amyloids have also been discovered, which are associated with the normal function of the host organism, termed “functional amyloid” (Maji et al., 2009), (Liebman and Chernoff, 2012), (Fowler et al., 2005), (Otzen, 2010). For example, yeast prions showed prion-like transmissive properties similar to human prion protein but provide a survival advantage to the host organism against harsh environmental conditions (Liebman and Chernoff, 2012), (Liebman and Chernoff, 2012), (Halfmann et al., 2012). Several mammalian functional amyloids have also been discovered where these amyloid support normal function rather than causing cell death (Barnhart and Chapman, 2006), (Maji et al., 2009), (Fowler and Kelly, 2012), (Chatterjee et al., 2022). Recently it was suggested that p53 aggregation and amyloid formation is associated with p53 loss- and gain-of-oncogenic function (Ghosh et al., 2017), (Navalkar et al., 2020b; Navalkar et al., 2021; Navalkar et al., 2022), (Silva et al., 2018), (Marques et al., 2022). These studies suggest that p53 amyloids could serve as an oncogene and may cause cancer initiation in cells (Ghosh et al., 2017; Navalkar et al., 2021; Navalkar et al., 2022), (Sengupta et al., 2022). Moreover, p53 amyloid formation not only leads to its loss of tumor suppressive function but might also cause the gain of tumorigenic function in cells by sequestering the other tumor suppressor proteins (Xu et al., 2011) and/or by prion-like p53 amyloid amplification (Forget et al., 2013), (Ghosh et al., 2017; Navalkar et al., 2021) similar to prion protein (Prusiner, 1998), (Aguzzi and Heppner, 2000).

In contrast to most of the neurodegenerative amyloid diseases (Chiti and Dobson, 2017), (Wolfe and Cyr, 2011), (Crews and Masliah, 2010), the p53 amyloid load in relation to the prognosis of cancer severity/grade is not been examined yet. To examine the relationship of p53 amyloid and cancer grades, we studied various grades of Indian patients’ biopsies of the stomach and oral cancers. We found that all the stomach and oral cancer tissues under studies contain p53 amyloids (**Fig. 1, Fig. 2 and Fig. 3**). Interestingly, we observed an increase in p53 amyloid with increased cancer grades for both cancer types (**Fig. 4**). This data support that widespread p53 deactivation and oncogenic gain of function of p53 could be associated with p53 amyloids. This is further supported by the fact that these p53 aggregates are neither functional nor able to bind cognate DNA sequence for their apoptotic activity supporting more cell survival in cancer similar to functional amyloid in yeast prion (Bradley et al., 2002; Edskes et al., 2014; Ness et al., 2002). Indeed, tumors containing p53 amyloids showed a striking reduction in the amount of p53 bound with response elements (genes responsible for apoptosis or cell cycle arrest such as p21, PIG, and Gadd45) (**Fig. 8A**) in higher cancer grades than in the lower grades, which is expected due to greater accumulation p53 amyloid load. This probably gives more advantages to the cell for cancer progression due to p53 amyloid formation (**Fig. 8K**). Previous data suggested that cancer associated mutations destabilized the p53 functional fold, which may induced aggregation and amyloid formation (Wang and Fersht, 2015b), (Moll et al., 1992; Moll et al., 1996), (Wang and Fersht, 2012), (Levy et al., 2011), (Rangel et al., 2014). Aggregation of wild-type p53 are also known in cancer cells and tissues (Moll et al., 1992; Moll et al., 1996), (Ostermeyer et al., 1996), (Wang and Fersht, 2015b). Consistent with this, the sequence information of the p53 gene in these cancer tissues showed p53 mutations in most of the cancer tissues and in few cancer tissues, we found wild-type p53 (**Fig. 5**). Interestingly, in contrast to previous study (Moll et al., 1995), our sequence data and p53 amyloid localization (using DAB staining) study showed that both wild-type and mutant p53 form either nuclear or cytoplasmic or nucleo-cytoplasmic p53 amyloids (**Fig. 6,7**). Irrespective of nuclear or cytoplasmic localization and/or mutations status (WT or mutant protein), p53 showed severe functionality loss in these cancer tissues suggests that misfolding of wild type or misfolding initiated by p53 mutation can produce p53 amyloids and subsequent loss of p53 function and gain of tumorigenic function of p53 (**Fig. 8**). In this context, a previous study showed a correlation between cytoplasmic p53 deposits and poor prognosis in high-grade serous ovarian carcinoma patients (Iwahashi et al., 2022). The study indicating that oncogenic cytoplasmic p53 aggregates can contribute to disease progression. In contrast, our study showed that the presence of p53 amyloid is correlated with cancer grades irrespective of the localization of these amyloids in nucleus and/or in cytoplasm suggesting that extent of p53 aggregation and amyloid formation could be more predictive for tumor prognosis in these cancers. Also there is possibility that p53 aggregation/amyloid formation either exclusively in nucleus or cytoplasm or both compartment could be prognosis factors depending upon the cancer types. The higher amount of p53 amyloid with increasing cancer grade suggests that similar to prion-like amplification/spread (Ness et al., 2002), (Chiti and Dobson, 2006), (Bradley et al., 2002)(Iwahashi et al., 2022), p53 amyloid amplification might result in widespread p53 inactivity and gain of tumorigenic function in the higher grade of cancers. We further asked whether, apart from self-amplification, the p53 amyloid might also sequester other p53 paralogs such as p63 and p73 (Xu et al., 2011), leading to a dominant negative effect as proposed for p53 mutations (Mantovani et al., 2019). Both p63 and p73 are known to display the transcriptional activity (Dötsch et al., 2010; Blandino and Dobbelstein, 2004) and tumor suppressor functions (McKeon, 2004). Therefore, p63/p73 sequestration by p53 amyloids might render them non-functional resulting in disease progression with increasing cancer grades. Indeed, we found increased colocalization of p63 and p73 isoforms with p53 amyloid in higher grades of both the cancers (**Fig. 6**). The data suggest that p53 amyloid formation, its prion-like amplification, and sequestration of other p53 paralogs might provide conducive environments for higher cancer grades. The present data, therefore, demonstrate that increased p53 amyloid formation can be a prognosis factor of cancer grade and p53 aggregation inhibitors (Soragni et al., 2016), (Palanikumar et al., 2021), (Ferretti et al., 2022) might be a valuable target against cancer.

## Material and Methods

### Chemicals and reagents

All the chemicals and reagents used for the study were of the highest purity and purchased from either Sigma-Aldrich (St. Louis, MO, USA) or Merck (Darmstadt, Germany). Double-distilled and de-ionized water was prepared using a Milli-Q system (Millipore Corp., Bedford, MA, USA). DNA extraction kit was obtained from Qiagen. The seprion ELISA kit was obtained from Microsens Biotechnologies (Cambridge, UK). The p53 activity ELISA kit was obtained from Cayman Chemicals (George Town, USA).

### Indian cancer patient’s tumor biopsies

A cohort of freshly frozen human oral and stomach cancer and their corresponding normal tissues were procured from the National Tumor Tissue Repository at the Tata Memorial Hospital, Mumbai, India. The entire study was approved by the Institutional ethics committee (IITB-IEC/2019/046), Indian Institute of Technology Bombay, Powai, Mumbai, India. The study included 60 tissues (54 tumor tissues and six control). These tissues were segregated into different grades. For oral cancer tissues, four grades (Grade I, II, III, and IV) were used, whereas, for stomach cancer tissues, Grade II and Grade III were used. Details of all the tissues are mentioned in **Table S1**. A minimum of 5 tissues per grade were selected for the study; however, in some grades, more than five tissues were used as per the prevalence. For the NGS study, a total of 48 tissues (44 tumors and four Normal) were taken to understand the p53 mutational status. For IHC studies, all 60 tissues were analyzed. The tissues were fixed and dehydrated, followed by clearing with xylene. The wax infiltration was carried out, and the tissues were sectioned (3-5 μM thickness) using a microtome and embedded onto glass slides.

### Hematoxylin and eosin (H&E) staining

The deparaffinization of the tissue sections was performed using the decreasing concentrations of xylene, starting with 100% xylene, followed by xylene and ethanol in a 1:1 ratio. This was further followed by rehydration in decreasing concentrations of ethanol from 100% to 50%, and finally washed with distilled water. Sections were then stained for 2 min with 0.5% hematoxylin solution. Subsequently, 0.8% eosin, prepared in 95% ethanol was used to stain the sections for 1 min, and the slides were kept in xylene for 1 hr. The sections were mounted using DPX (Dibutyl-phthalate Polystyrene Xylene) mounting media and observed under a Leica DMi8 (Leica Microsystems, Germany) fluorescence microscope fitted with Andor Zyla cCMOS camera (Oxford Instruments, UK).

### Immunohistochemistry of tissues

Paraffin-embedded fixed tumors and the corresponding normal tissue sections were used for the immunohistochemistry study. The entire work plan and protocols were approved in advance by the Institutional ethics committee (IITB-IEC/2019/046), Indian Institute of Technology Bombay, Mumbai, India. Tissues were deparaffinized and rehydrated, as mentioned above. The enzymatic antigen retrieval was performed by incubating the sections for 2 min with TrypLE™ Express Enzyme (ThermoFisher) at 37 °C. The sections were then washed with Tris-buffered saline (TBST) with 0.1% tween-20, pH 7.4) and then incubated with 0.2% Triton X-100 in TBST for 10 min. The section blocking was done with 2% BSA in TBST. The sections were incubated overnight at 4 °C with primary antibodies, such as mouse monoclonal anti-human p53 protein DO-1 (1:200) (Santa Cruz Biotechnology, Dallas, TX, USA) or anti-human pab240 (1:500) (Santa Cruz Biotechnology, Dallas, TX, USA) and rabbit polyclonal oligomer-specific (A11)(Kayed et al., 2003) (1:500) or amyloid-specific (OC) (1:500) antibody (Kayed et al., 2007). Tissues were washed with TBST followed by incubation with the secondary antibody such as anti-mouse FITC-488 (1:1000) or goat anti-rabbit Alexa Fluor-647 (Life Technologies, Thermo Scientific, USA) at room temperature for 2 h. To study the coaggregation of p53 with p63 or p73, rabbit monoclonal anti-human p53 protein SP5 (1:200) (Invitrogen, USA) and mouse monoclonal anti-p63 antibody (1:500) or anti p73 antibody (1:500) (Santa Cruz Biotechnology, Dallas, TX, USA) The Thioflavin S (Thio S) staining was performed after immunostaining with an anti-p53 primary antibody and subsequent incubation with an anti-mouse Alexa Fluor-555-conjugated secondary antibody (1:1000 dilution). After antibody staining, the sections were stained for 2 min with filtered 0.6% Thio S solution (Sigma-Aldrich) in the dark. The sections were washed twice with initially 50% ethanol and then with TBST buffer. The sections were then mounted with 1% DABCO (1,4-diazabicyclo-[2.2.2] octane, Sigma-Aldrich) prepared in 90% glycerol and 10% phosphate-buffered saline (PBS) and left for drying. Imaging was performed using Zeiss Axio Observer.Z1 inverted confocal fluorescence microscope (Zeiss, Germany) fitted with a high-speed microlens-enhanced Nipkow spinning disc (CSU-X1, Yokogawa Electric Corporation, Tokyo, Japan).

### Isolation of the tissue amyloid fibrils

The total pool of amyloid fibrils from oral and stomach tumor tissues was isolated using previously reported methods (Haltia et al., 1990) with certain modifications (Ghosh et al., 2017). Tumor tissues (150–200 mg) were homogenized for 20 min in 500 μl of 0.15 M NaCl and centrifuged at 9000 × g for 1 h at 4 °C. The supernatant is discarded, and the pellet was re-homogenized in the same buffer and centrifugation was done at 9000 × g for 1 h at 4 °C. The supernatant was discarded, and the resulting pellet was homogenized in 500 μl of 0.05 M Tris-HCl, 3 mM NaN3, 0.01 mM CaCl2, pH 7.5. Collagenase type I (Himedia) was added to the weight of the total tissue at a ratio of 1:100 and incubated at 37 °C overnight. The next day, the homogenate was centrifuged at 28000 × g for 1 h at 4 °C using the ultracentrifuge. The resulting pellet was then homogenized in 300 μl of 0.15 M NaCl and centrifuged at 28000 × g for 1 h at 4 °C. This step was repeated many times until the absorbance of the supernatant reached below 0.3. Further, the resulting pellet was then suspended in 200 μl distilled water followed by homogenization which was then centrifuged for 1 h at 28000 × g at 4 °C. All supernatants were then pooled together and NaCl (0.15 M) was added for the precipitation of fibrils. This mixture was centrifuged for 1 h at 28 000 × g at 4 °C. The final pellet containing the total amyloid fibrils was suspended in PBS and stored at 4 °C until use.

### Transmission electron microscopy

10 μl of the isolated amyloid fibril obtained as mentioned above were spotted on to a copper-coated formvar grids (Electron Microscopy Sciences, Hatfield, PA, USA). The grid was washed with MQ water and stained for 20 min with 10 μl 0.1% uranyl formate solution (Electron Microscopy Sciences). The uranyl formate solution was prepared freshly and filtered with a 0.22 μm sterile syringe filter (Millipore, Billerica, MA, USA) before use. For immunoelectron microscopy, 10 μl anti-p53 DO-1 antibody (1:10) was spotted on the grid carrying the samples for 20 min. The grid was subsequently washed with MQ water followed by incubation for 20 min with 10 μl of anti-mouse 10 nm gold-labeled secondary antibody (1:10) (Sigma-Aldrich). Further, the grid was again washed and stained with 0.1% uranyl formate solution. The images were acquired at X10000 magnifications at 200 kV using JOEL FEG-TEM 200 (JEM-2100 F) (JEOL, Tokyo, Japan). Recording of images was done digitally using the Gatan Microscopy Suite® (Gatan, USA). For Co-Immuno TEM, 10 μl anti-p53 (SP5, Invitrogen) antibody (1:10) and 10 μl anti-p63 antibody (Santacruz) was spotted on the grid carrying the samples for 20 min. The grid was subsequently washed with MQ water followed by incubation for 20 min with 10 μl of anti-mouse 10 nm gold-labelled secondary antibody (1:10) (Sigma-Aldrich) and 10 μl of anti-rabbit 5 nm gold-labelled secondary antibody (1:10) (Sigma-Aldrich). The grid was washed, stained with 0.1% uranyl formate solution and acquire images at X10000 magnifications at 200 kV using JOEL FEG-TEM 200 (JEM-2100 F) (JEOL, Tokyo, Japan). Recording of images was done digitally using the Gatan Microscopy Suite® (Gatan, USA).

### FTIR imaging

Fixed tissue sections were used for the FTIR Imaging. Frozen tissue sections were deparaffinized and rehydrated as mentioned above. The tissue samples were scrapped and kept on the BaF2 window (38 × 19 × 4 nm) (Technosearch Instruments, India). The BaF2 is known for its low absorbance in the whole UV-IR wavelength spectrum (200 nm to 12 µM) and also provides resistance in high-energy radiation. For background correction, a clean area of each BaF2 slide was measured and subsequently subtracted from sample measurements. During FTIR spectra acquisition, a Vertex-80v vacuum optics bench (Bruker, Germany) was used, and for recording the FTIR spectra, a Vertex-80 FTIR machine (Bruker, Germany) was attached with a 3000 Hyperion microscope. The tissues were imaged at 15X magnification (2.7 µm resolution) in the focal plane array (FPA) mode in the wavenumber range of 1600-1700 cm^-1^ corresponding to amide-I stretching (C=O) frequency of peptide bond. For the analysis of FTIR spectra, the OPUS-65 v6.5 software was used. Individual points on the tissue images were selected for the analysis. To eliminate any contribution from water in the FTIR spectrum near 1650 cm^-1^, background correction was done. The spectra were then subjected to baseline correction followed by Fourier Self Deconvolution (FSD) using the Lorentzian deconvolution function in the wavenumber range of 1700-1600 cm^-1^. Briefly, the deconvolution technique was used to get the single sharp peaks from the convoluted or broadened spectra. Notably, the deconvolution was performed by feeding noise and band deconvolution factors optimized for minimum noise and maximum sharpness in the spectra. After that, the peaks were assigned according to the protein secondary structures and were best fitted by the auto-fitting method in the OPUS 65 software with minimum root mean square (RMS) error. The area under each resulting peak, the fractional contribution of individual secondary structures, was integrated by the peak integration method in the OPUS-65 software. The integration value of the β-sheet is divided from the sum of all the secondary structures and multiplied by 100 to obtain the % abundance of β-sheet in every FTIR spectrum. FTIR imaging and double immunofluorescence with anti-p53 antibody along with OC antibody were performed on the adjacent tissue sections of the same slides.

### Dot blot assay

All the different grades of oral and stomach cancer tissues were lysed and homogenized using RIPA buffer, supplemented with the Protease Inhibitor cocktail (PIC). The lysate was centrifuged at 3000 g for 5 min at 4^ο^ C to remove the tissue debris. The tissue lysate or amyloid fraction containing p53 protein was used for dot blot assay. For the detection of p53 amyloid, 20 μg of lysate was used. 4 ul of the samples were spotted directly on the nitrocellulose membrane. After drying, the membrane was blocked using blocking solution of 5% non-fat skimmed milk powder (Himedia, India) prepared in TBST for 1 h. Further, the membrane was washed with TBST and incubated overnight at 4°C with anti-p53 antibody, 1:200 dilution or OC, 1:500 dilution. After incubation, the membranes were washed thrice with TBST for 10 min, followed by 2 h incubation at RT with anti-mouse HRP tagged secondary antibody for p53 and anti-rabbit for OC (ThermoFisher Scientific, USA) with 1:10000 dilution. Nonspecific binding was removed by washing the blot three times with TBST for 10 min. The protein signals were developed with SuperSignal West Femto kit (Thermo Scientific).

### Next Generation Sequencing

A total of 48 oral and stomach tissues (44 tumor tissues and 4 normal tissues), as mentioned in **Fig. S1,** were taken for the Next Generation Sequencing of p53 gene. The genomic DNA was isolated from these tissues using QIA amp DNA Mini Kit (Qiagen) based on the manufacturer’s instructions. The gDNA quality and quantity were assessed using Nanodrop 2000 and Qubit (Thermo Scientific, USA), respectively. The sequencing library was prepared using an Illumina-compatible Accel Amplicon library prep kit and TP53 Comprehensive Panel (Swift Biosciences) at Genotypic Technology Pvt. Ltd., Bangalore, India. Briefly, 20 ng of Qubit-quantified genomic DNA was taken as template for Multiplex PCR using Reagent G1 of the TP53 Comprehensive Panel, and PCR amplification was carried out for 22 cycles, following the manufacturer’s instructions. The amplicons were bead-purified in a 1.2X bead: sample ratio, followed by Indexing whereby unique combinations of dual indices were added to the amplicons. Finally, the barcoded samples were bead-purified in a 0.8X bead: sample ratio. The purified libraries were quantified using Qubit fluorometer (Thermo Fisher Scientific, MA, USA) and qPCR assays. The libraries were paired-end sequenced using Illumina HiSeq X Ten sequencer (Illumina, San Diego, USA) for 150 cycles following the manufacturer’s instructions.

Raw reads obtained from Illumina HiSeq X Ten sequencing for 48 samples were processed using Trim Galore-v0.4.01to generate high-quality reads by removing adapters, reads with less than Q30 quality score, and reads with length less than 20. Next, these processed reads were mapped against chromosome 17 of the Grch37 genome using Bowtie2 v2.2.5 2 aligner to generate alignment files. The alignment files were used for the removal of PCR duplication and adding read group information using Picard v1.102 3 tool followed by GATK v4.1.4.1 4 for performing Realigner Target Creator, Indel Realigner and Haplotype Caller to generate variants. Variants were generated for 44 samples (out of 48 samples) in GATK Haplotype Caller by using targeted regions bed file and reference (chromosome 17) sequence. Finally, the variant files were used for annotation using Variant Studio v3.0 5 tool. The sum of the mutant frequency was calculated by dividing the sum of total alternate frequency obtained for each patient by the number of tissues sequenced for that specific grade.

### Seprion-ELISA Amyloid quantification Assay

The tumor tissue was weighed equally (∼ 30 mg), and ice-cold NP-40 buffer, including a protease inhibitor cocktail (complete inhibitor, Roche), was added to prepare a 2.5% (w/v) lysate. Tissues were homogenized and incubated on ice for 30 min. The p53 amyloid load was determined using the Seprion-ELISA (Microsens) as described earlier (Haltia et al., 1990). The assay is based on a polyionic, high-molecular-weight ligand coated to the ELISA plate’s surface. Only the aggregated and/or amyloid proteins can bind to the ligand in the presence of seprion capture buffer. Further, anti-p53 antibody was used for the sandwich ELISA to detect only the p53 aggregates/fibrils. The tissue lysate was added to the assay plate, and absorbance was measured using a plate reader. The absorbance is proportional to the bound amount of aggregated/fibril p53.

### Dab Staining

The Fixed tumor and normal tissues were deparaffinized rehydrated and enzymatic antigen retrieval was performed, as mentioned earlier. The endogenous peroxidase was quenched by incubating slides with 3% H2O2 for 15 min. The tissue sections were subsequently washed with Tris-buffered saline with 0.1% tween-20 (TBST), pH 7.4, which was subsequently treated with 0.2% Triton X-100 in TBST for 10 min. We used a blocking buffer of TBST containing 5% BSA to block nonspecific antigenic sites. The sections were then incubated with mouse monoclonal anti-human pab240 primary antibody (Santa Cruz Biotechnology, Dallas, TX, USA) (1:500) overnight at 4 °C. Tissues were further incubated for 2 h at room temperature with HRP-tagged secondary antibody of goat anti-mouse (1:1000) (Life Technologies, Thermo Scientific, USA). The sections were then washed with TBST three times. The sections were then incubated with dab solution Sigma-Aldrich (St. Louis, MO, USA) with 0.2% NiCl2 for 20 min and counterstained with haematoxylin. The sections were dehydrated again with increasing concentrations of ethanol and xylene. The slides were mounted with DPX (Dibutyl-phthalate Polystyrene Xylene) mountant and observed using Leica DMi8 (Leica Microsystems, Germany) fluorescence microscope fitted with Andor Zyla cCMOS camera (Oxford Instruments, UK), and images were analyzed using ImageJ2 software. The dab intensity was quantified to determine the nuclear and cytoplasmic misfolded p53 in the sections. For that, the color deconvolution plugin of Image J was used. The built-in stain vector Haematoxylin and DAB (H DAB) was selected. Images stained only with DAB were selected, and total intensity was measured. Further, the threshold was adjusted so that only the nuclear intensity could be measured. The nuclear Dab intensity was then subtracted from the total intensity to obtain the cytoplasmic Dab intensity in the sections.

### Chromatin Immunoprecipitation

Chromatin immunoprecipitation assay was carried out to assess the functional status of p53 using Magna ChIP™ G Tissue Kit (Millipore, USA). Briefly, 30 mg of fresh tissue biopsy was weighed and washed with PBS to remove any attached impurity. 1% formaldehyde was added for crosslinking, followed by glycine to stop the reaction. The tissue was manually disrupted, followed by sonication for 5 cycles with 1 min on and off, and centrifuged at 4 °C to remove any cell debris. The supernatant was removed to fresh microfuge tubes in 125 µl aliquots. The supernatant was diluted in dilutant buffer and was incubated overnight in the presence of 20 µl of fully resuspended protein G magnetic beads and 10 µl anti-p53 antibody. The Protein G magnetic beads were pelleted with the magnetic separator rack, and the supernatant was removed completely. The Protein G bead-antibody/chromatin complex was washed by resuspending the beads in 0.5 ml of each of the cold buffers in the order as mentioned (Low Salt Immune Complex Wash Buffer, High Salt Immune Complex Wash Buffer, LiCl Immune Complex Wash Buffer, one wash, TE Buffer, one wash and incubating for 3-5 minutes on a rotating platform followed by magnetic clearance and careful removal of the supernatant fractions. Protein-DNA crosslink reversal was carried out, and DNA was eluted and purified using QIAquick PCR purification kit (QIAGEN, Valencia, CA, USA) according to the manufacturer’s instructions. The qPCR was performed in 20 μl SYBR Green reaction mixture using a Real-time PCR system (Agilent AriaMx Real-time PCR System, Agilent, USA). The ChIP experiments and qPCR were performed in triplicates. The enrichment/input values were calculated as: ΔCT=CT(ChIP)−[CT(Input)−LogE (Input dilution factor)] where E is the specific primer efficiency value; % Enrichment/Input=E−ΔCT. All the primers used in this study are listed in **Table S2**.

### Immunoprecipitation

The different grades of oral and stomach tissues were suspended in RIPA lysis buffer (20 mM Tris-HCl, pH 8.0, 137 mM NaCl, 1% NP-40, 2 mM EDTA) with protease inhibitor cocktail (Roche, Switzerland) and manually homogenized on ice. The mixture was centrifuged at 8 000 rpm at 4 °C for 20 min. The obtained supernatant was incubated with the anti-p53 antibody (SP5, Invitrogen) overnight at 4 °C under rotation. Next day, 100 μl of Sepharose G beads (Thermo Fisher Scientific, MA, USA) were added to the solution and incubated at 4 °C under constant agitation for 4 hrs. The solution was then centrifuged at 3000 rpm for 1 min to precipitate the beads. The supernatant was discarded and the elution was performed by mixing the equal amount of SDS loading dye to the beads and heating the samples at 95 °C for 10 min. The eluted fractions were loaded in the SDS gel and western blot was performed with anti-mouse p63 and p73 primary antibodies (Santa Cruz Biotechnology, Dallas, TX, USA)(1:500) as mentioned above. All the original uncropped western blot images are added as a separate file (Fig S8F-I).

### Database analysis

The TCGA data and its specific grade information were retrieved from UCSC Xena (https://xena.ucsc.edu/). We previously established the p53 amyloid-specific alteration of gene expressions in cells (Navalkar et al., 2021; Navalkar et al., 2022). Utilizing those unique gene-sets, we examined the single-sample gene set enrichment analysis (ssGSEA) (Subramanian et al., 2005) using python package gseapy (https://github.com/zqfang/gseapy). This quantifies the amyloid-specific uniquely differential expressed genes with the grades of various cancer types in the TCGA database. R version 4.2.0 was used for all statistical analysis and the plots were generated using ggplot2 function.

### Statistical Analysis

The statistical significance was calculated using an unpaired two-tailed t test or one-way ANOVA followed by Tukey’s multiple Comparison test. All the data presented here are the mean ± standard error. At least three biologically independent experiments were performed unless otherwise stated in the figure legends. The p-value for the significance is *p < 0.05, **p < 0.01, ***p < 0.001; non-significant (NS p > 0.05). Graphpad prism was used for calculating the statistical significance.

## Acknowledgments

We acknowledge the National Tumor Tissue Repository (NTTR), Indian Council for Medical Research (ICMR) at the Tata Memorial Hospital (TMH), Mumbai, India, for providing human cancer tissue; Dr. Omshree Shetty, Tata Memorial Hospital, Dr. Sushil Kumar, IIT Bombay for providing valuable inputs, Mr. Pradeep Kadu for making the schematic and Ms. Manisha Poudyal for her inputs in figure re-arrangements. We also acknowledge the Center for Research in Nano Technology and Science (CRNTS) and Industrial Research and Consultancy Centre (IRCC), IIT Bombay, for FTIR, electron microscopy, Confocal facility and the Department of Biotechnology (DBT) (BT/PR9797/NNT/28/774/2014) Government of India, Wadhwani research center for Bioengineering (WRCB), the Department of Science and Technology (DST), Ministry of Science and Technology, India (EMR/2014/001233 and CRG/2019/001133) for financial support. The authors wish to acknowledge the DBT/Wellcome Trust India Alliance Fellowship [IA/E/17/1/503663] awarded to Shinjinee Sengupta for financial support. Ajoy Paul acknowledges the Council of Scientific and Industrial Research (CSIR), Government of India, for the Shyama Prasad Mukherjee fellowship (SPMF).

## Author Contributions

S.S, N.S, A.P, D.D, D.C, S.M, L.G, and J.D performed the experiment. S.S, N.S, A.P, D.D, D.C, S.M, L.G, J.D, and Y.M analyzed the data. The study was conceived by S.K.M and designed by S.S, S.K.M, and M.K.J. S.S, N.S, A.P. D.D, and D.C participated in the manuscript writing. S.S and S.M prepared figures and illustrations.

## Data availability statement

The authors state that all the data supporting the findings of this study are reported within the paper and in the supplementary information files. All the data analysis was performed using published tools and has been cited in the paper and in the supplementary file.

