## Supplementary File for "p53 amyloid pathology with cancer grades and p53 mutations"

**This file includes:**

Supplementary Figures S1 to S8

Supplementary Tables S1 and S2

**Supplementary Figures:**

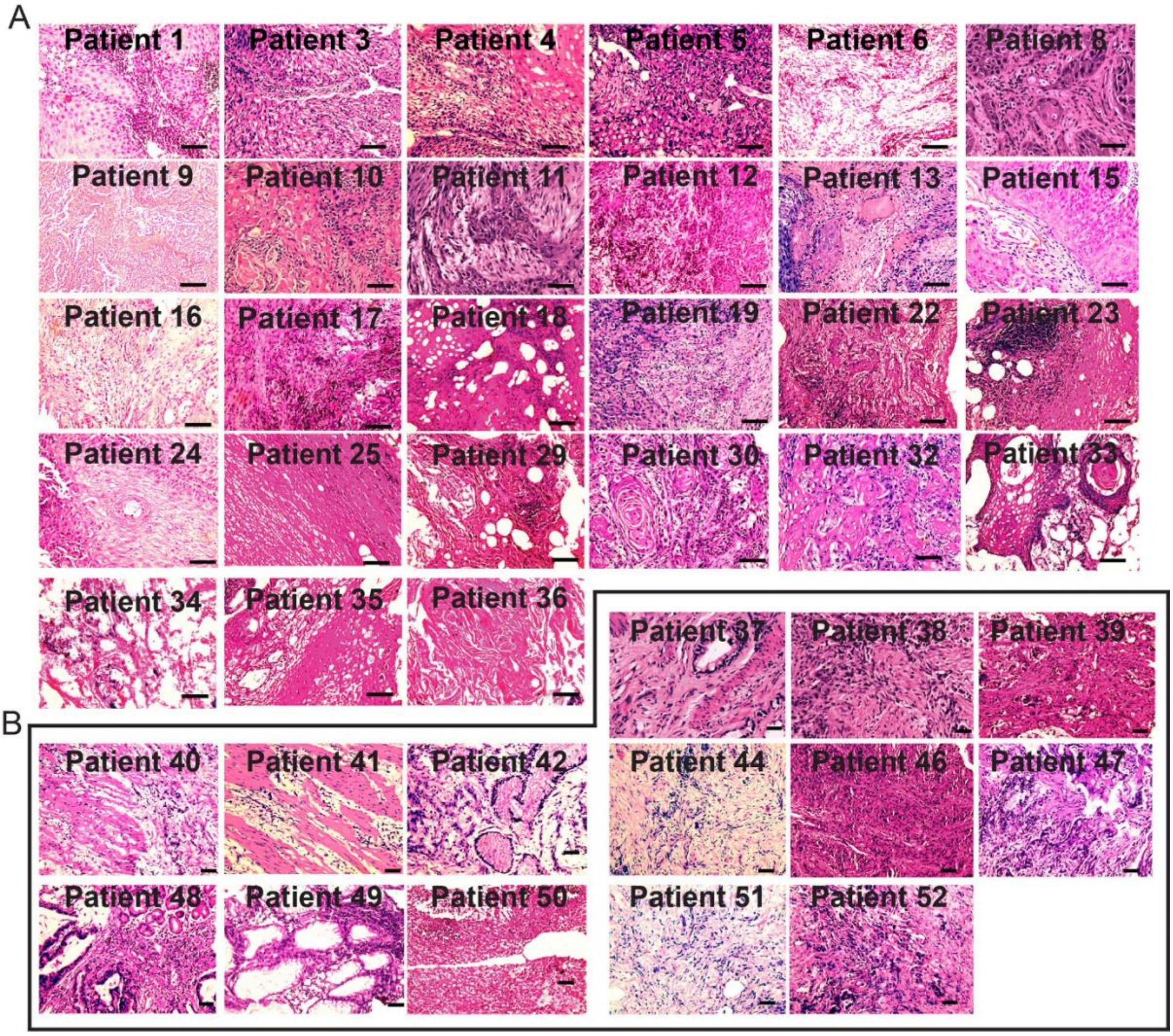


***Fig. S1. Hematoxylin and eosin staining of oral and stomach cancer tissues.*** Various cancer tissue sections from (a) oral and (b) stomach origin from different patient biopsies were deparaffinized, hydrated and stained with hematoxylin and eosin. The sections were examined using microscope with 20x magnification. The representative patient's oral cancer and stomach cancer tissues (marked with black box) of different grades showing differential staining with hematoxylin and eosin. The biopsies showed the presence of large number of nuclei, which were highly stained with hematoxylin. Scale bars are 50 μm. Images are representative of 3 independent experiments.

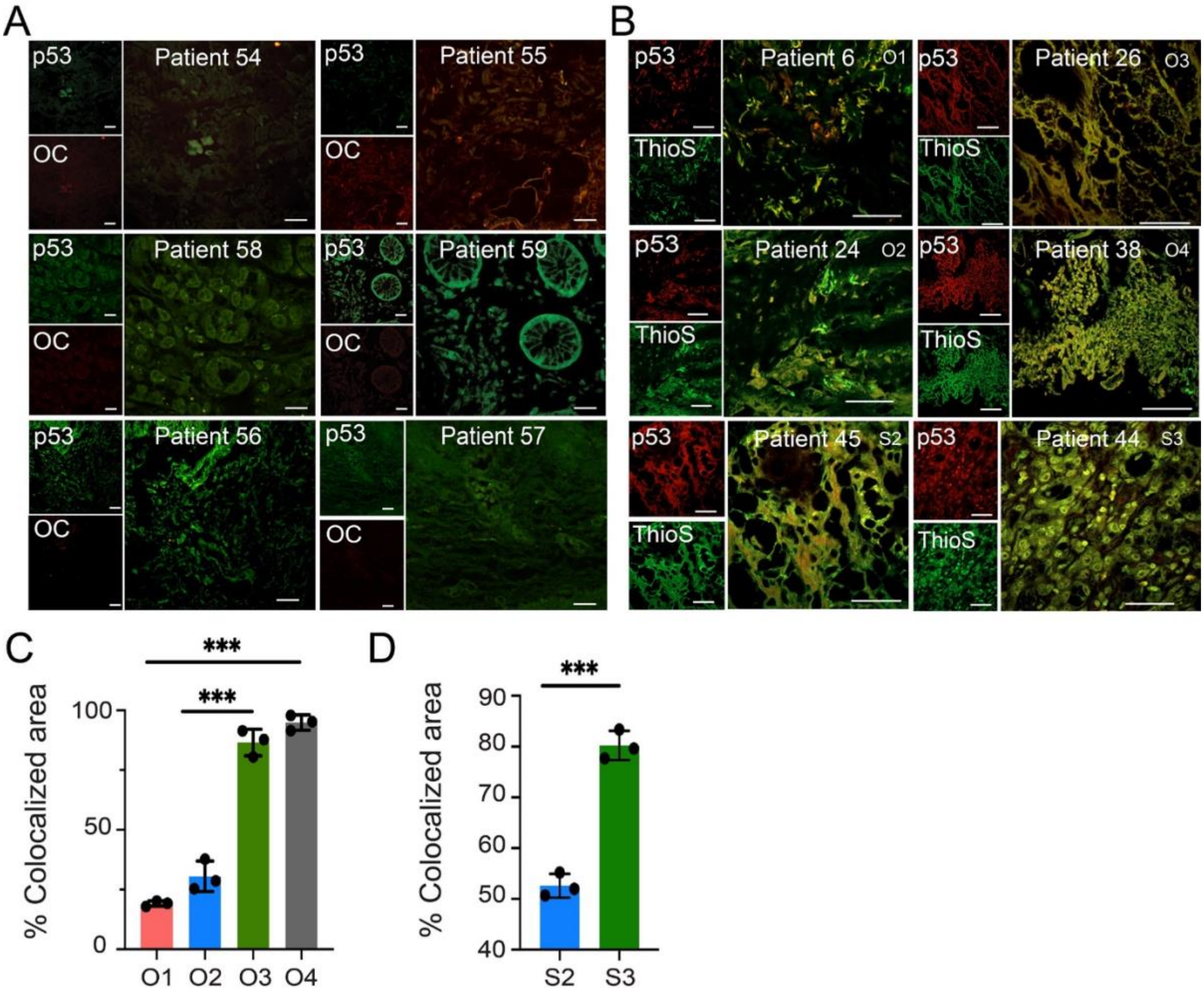


***Fig. S2. p53 with OC and ThioS staining of oral and stomach tissues.*** ***a.*** Immunofluorescence study of p53 in human oral and stomach noncancerous (normal) tissues using anti-p53 antibody (DO-1, 1:500 dilution) and amyloid specific antibody, OC (Abcam, 1:500 dilution) showing no significant co-localization of p53 and OC signals. The data suggesting the absence of p53 amyloids in normal tissue. b. Immunofluorescence study of p53 amyloids in oral and stomach cancer tissues using anti-p53 antibody (DO-1, 1:500 dilution) and amyloid-specific dye, ThioS. The co-localization of p53 and ThioS signals was observed in tumor tissues. Scale bars are 50 µm. Images are representative of 3 independent experiments. (c,d) Quantification of percent colocalized areas of p53 with ThioS dye showing the amyloid load in different cancer grades for both oral (c) and stomach (d). Representative images are shown although 3 patient tissues from each cancer grade is analysed. The values were plotted as mean ± s.e.m., n=3 independent experiments.

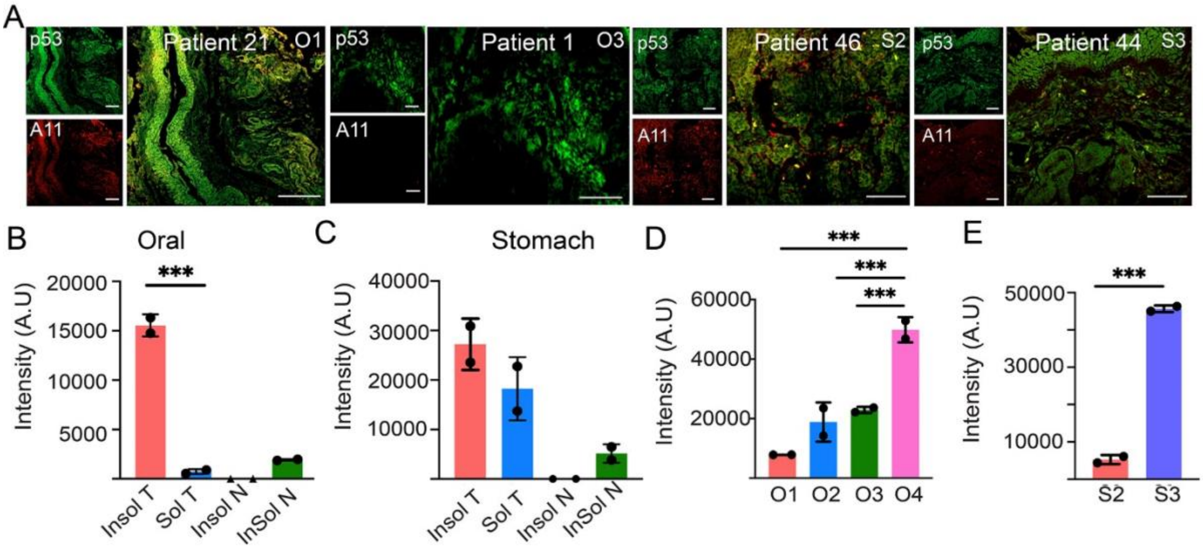


***Fig. S3. p53 amyloids and oligomeric status in cancer tissues.*** a. Double immunofluorescence study using anti-p53 antibody (DO-1) and oligomer specific antibody, A11 (Abcam, Cambridge, United Kingdom) in oral and stomach cancer tissues. The co-localization of p53 and A11 signals indicating the presence of p53 oligomers in lower grades of tumor tissue. Negligible colocalization was observed in higher grades of cancer tissues in both the cancer types. Scale bars are 50 µm. Images are representative of 2 independent experiment. b. Soluble and insoluble fractions of tissue lysate from higher cancer grades (Oral Grade III & Stomach Grade III) were separated using ultracentrifugation, and western blot analysis was performed to analyze the p53 status (Figure 4e). Quantification of p53 band intensity using Image J for both oral (b) and stomach (c). In both the cancer types, p53 was observed to be accumulated in the insoluble fraction (Insol T represents the insoluble fraction of the tumor tissue lysate, Sol T represents the soluble fraction of the tumor tissue lysate, Insol N represents the insoluble fraction of the normal tissue lysate and Sol N represents the soluble fraction of the normal tissue lysate). Negligible p53 expression was seen in the normal tissues. (d, e) Quantification of p53 amyloid load using OC antibody as depicted in the dot blot (Figure 4f) using Image J in different cancer grades for both oral (d) and stomach (e). The tissue lysate was quantified using Bradford and equal volume of the tissue lysate was spotted on the nitrocellulose membrane. The p53 amyloid load was observed to increase significantly with increase in the cancer grade for both the cancer types. The values were plotted as mean ± s.e.m., n=2 independent experiments.

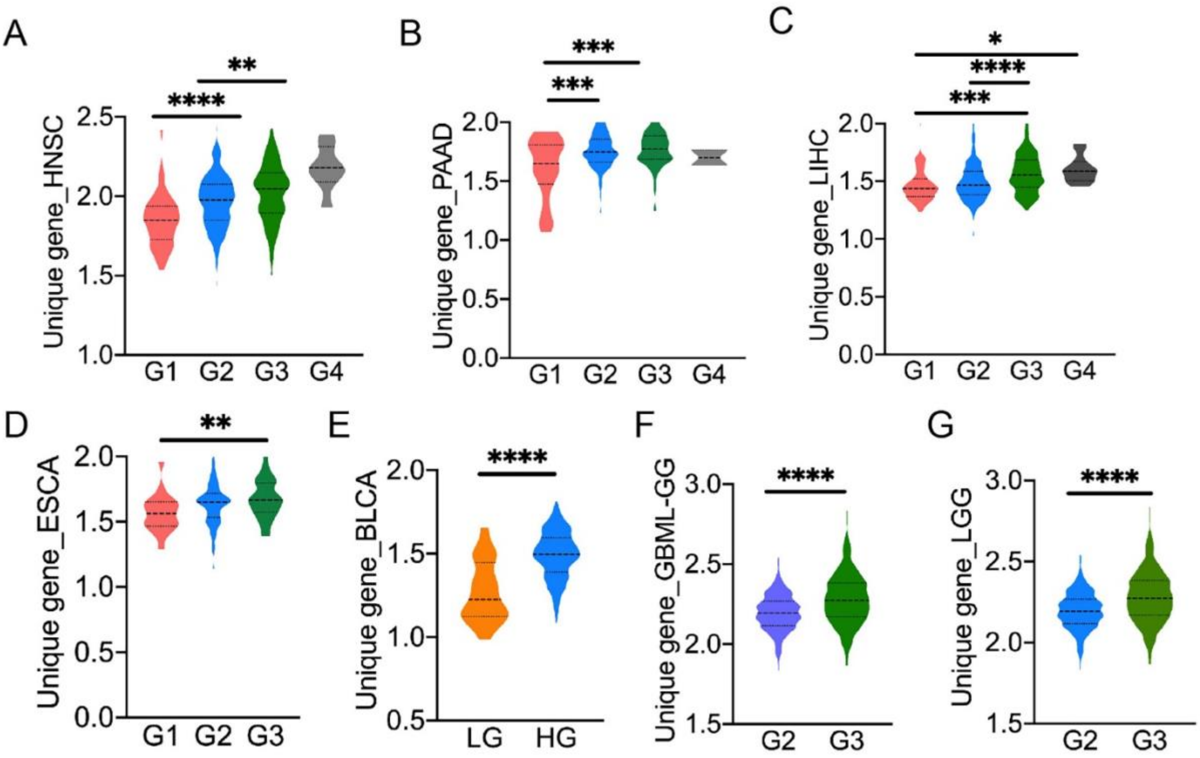


**Fig S4.** ***Cancer grade-based gene expression profile of the "p53-amyloid specific unique genes" in various cancer types using TCGA database.*** From the previous studies[1,2], we identified the differentially expressed unique genes associated with p53 amyloid formation using microarray. These genes were examined with the TCGA database, which showed higher alteration of gene expressions in higher grades of various cancers a. Head and Neck Cancer (HNSC), b. Pancreatic adenocarcinoma (PAAD), c. Liver Hepatocellular Carcinoma (LIHC), d. Esophageal Cancer (ESCA) e. Bladder Cancer (BLCA), f. Glioblastoma multiforme (GBM) and g. Low-grade gliomas (LGG). Statistical significance is (P<0.05) shown.

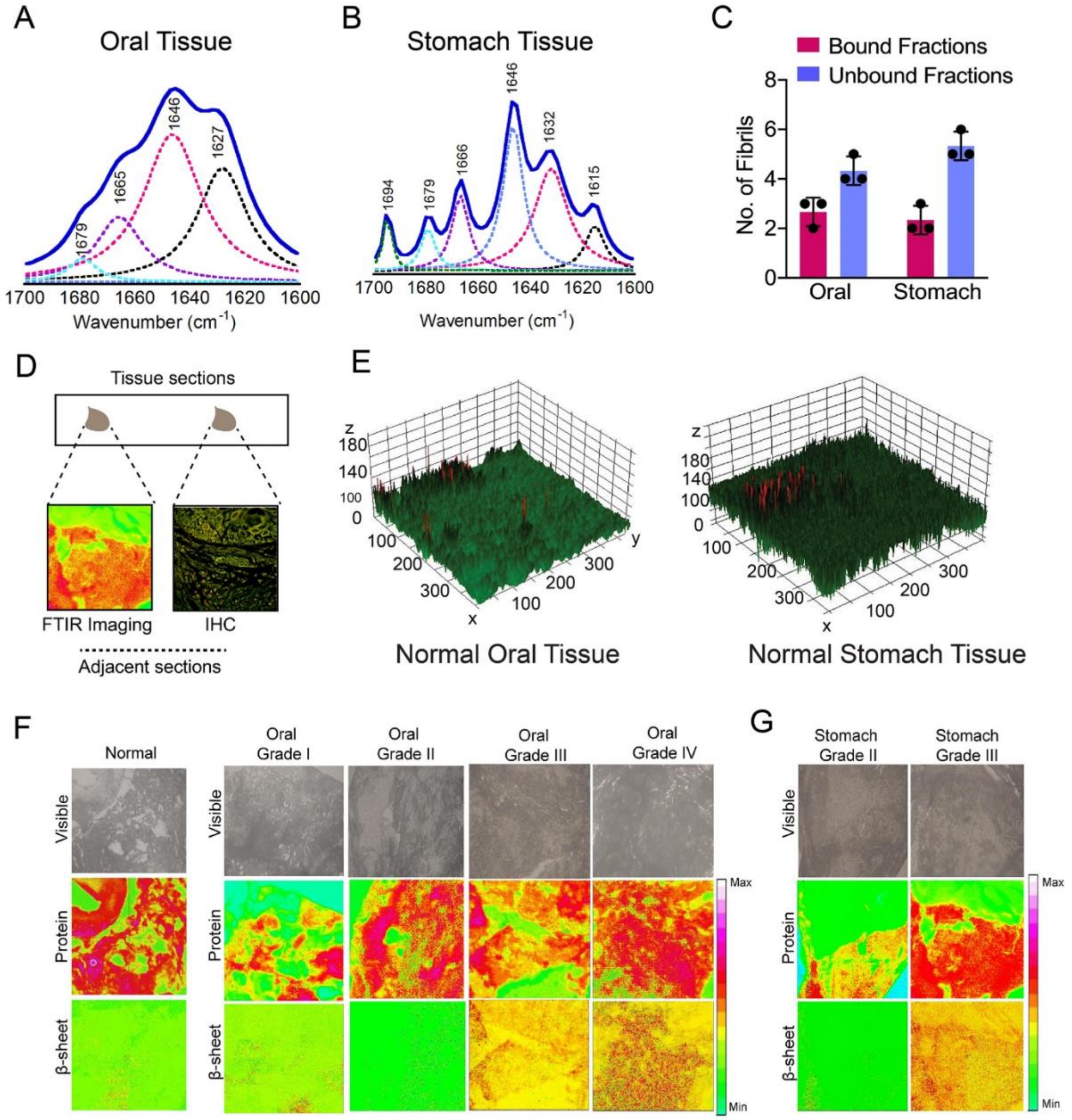


***Fig. S5. Biophysical characterization of p53 amyloid isolated from cancer tissues and FTIR imaging of oral and stomach tissues.*** FTIR spectra of amyloid fibrils isolated from oral (a) and stomach cancer tissues (b) in the amide I region (1600–1700 cm−1). Amyloid fibrils isolated from oral tissues showing FTIR peaks at 1627 in the amide-I region, indicating the presence of β-sheet secondary structure. Amyloid fibrils isolated from stomach cancer tissues showing FTIR peaks at 1615 and 1632 cm−1, which indicates the presence of β-sheet structure. (c) The population of bound and unbound fraction of the tissue isolated fibril population of the 10 nm gold-labeled secondary antibody against p53 antibody in oral and stomach biopsies. Values represent mean ± SEM for n=3 independent experiments. d. FTIR imaging of oral and stomach tissue sections of various grades. Schematic showing the tissue sections taken for FTIR imaging and corresponding immunostaining from the adjacent tissue section of the same tissue slide. (e). 3D surface plot showing the amyloid load of normal tissues oral (left panel) and stomach (right panel) analysed from FTIR imaging. (f). Top panel showing white light images. The middle panel shows the overall protein pool in the respective section with all possible secondary structures of the proteins (amide I C=O stretching: 1700–1600 $cm^{-1}$). The bottom panel showing the β-sheet rich amyloid region (1640–1620 $cm^{-1}$) of the corresponding protein pool, present in the respective region of the normal tissue (left), oral tumor tissues (middle) and stomach (right) tumor tissues. g. FTIR imaging of stomach cancer tissues showing β-sheet rich amyloid region.

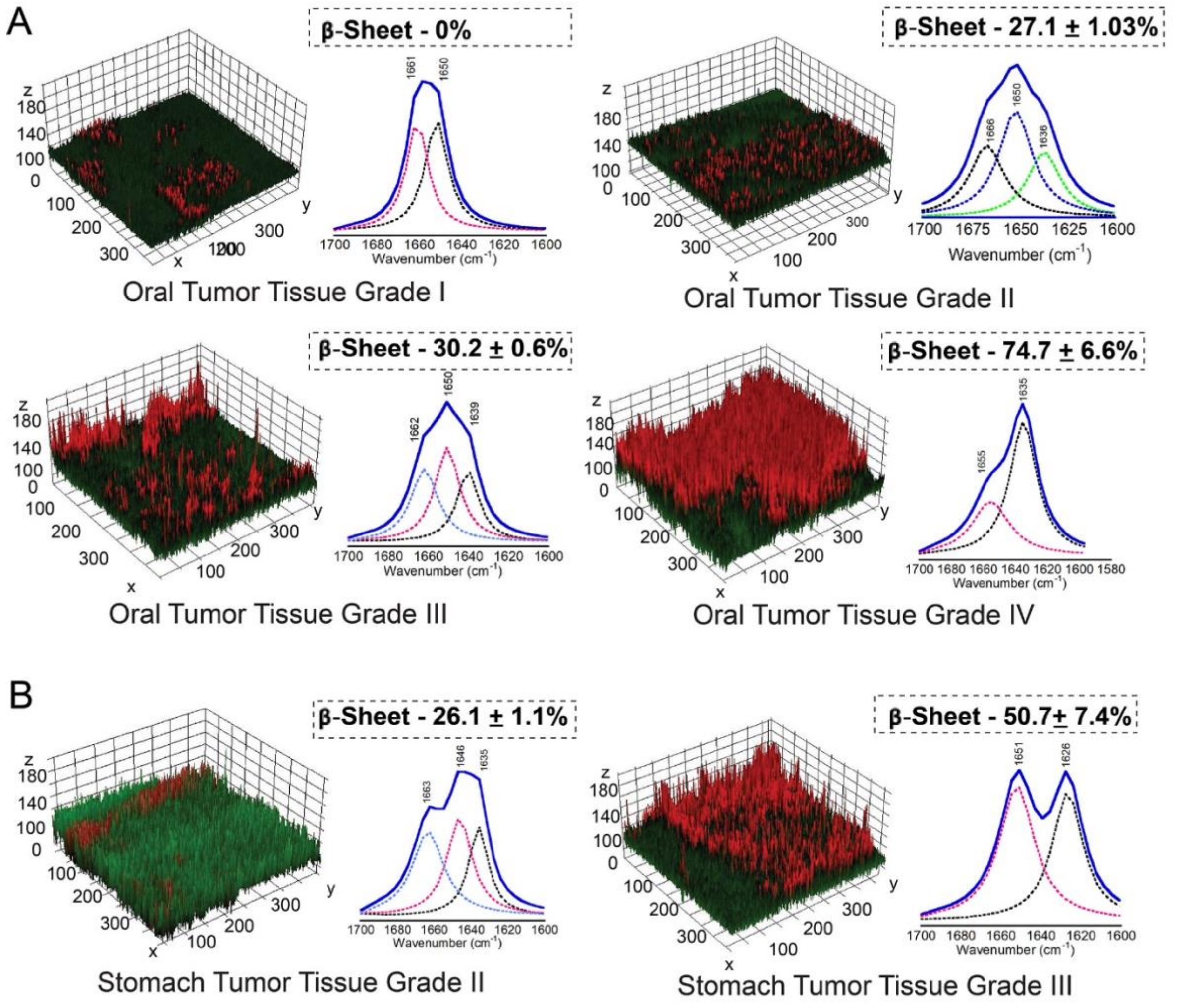


***Fig. S6. 3D surface plot of amyloid load from FTIR imaging of Oral and stomach tissues.*** FTIR spectroscopic imaging was performed for oral and stomach tissue sections of various grades. 3D surface plot showing the amyloid load (β-sheet rich amyloid region (1640–1620 $cm^{-1}$) analysed from FTIR imaging of Oral (a) and stomach (b) tissues. The corresponding FTIR signature is shown for each of the oral/stomach tumor tissue grade. Interestingly, the β-sheet content (1640-1620 $cm^{-1}$) is absent in oral tumor tissue grade I, but the quantity is enhanced gradually for the oral tumor tissue grade II, III and IV. The similar trend is observed for the stomach tumor tissue grade II and III, where stomach tumor tissue grade III possesses significantly higher β-sheet content than the stomach tumor tissue grade II. The values were plotted as mean ± s.e.m., n=2 independent experiments.

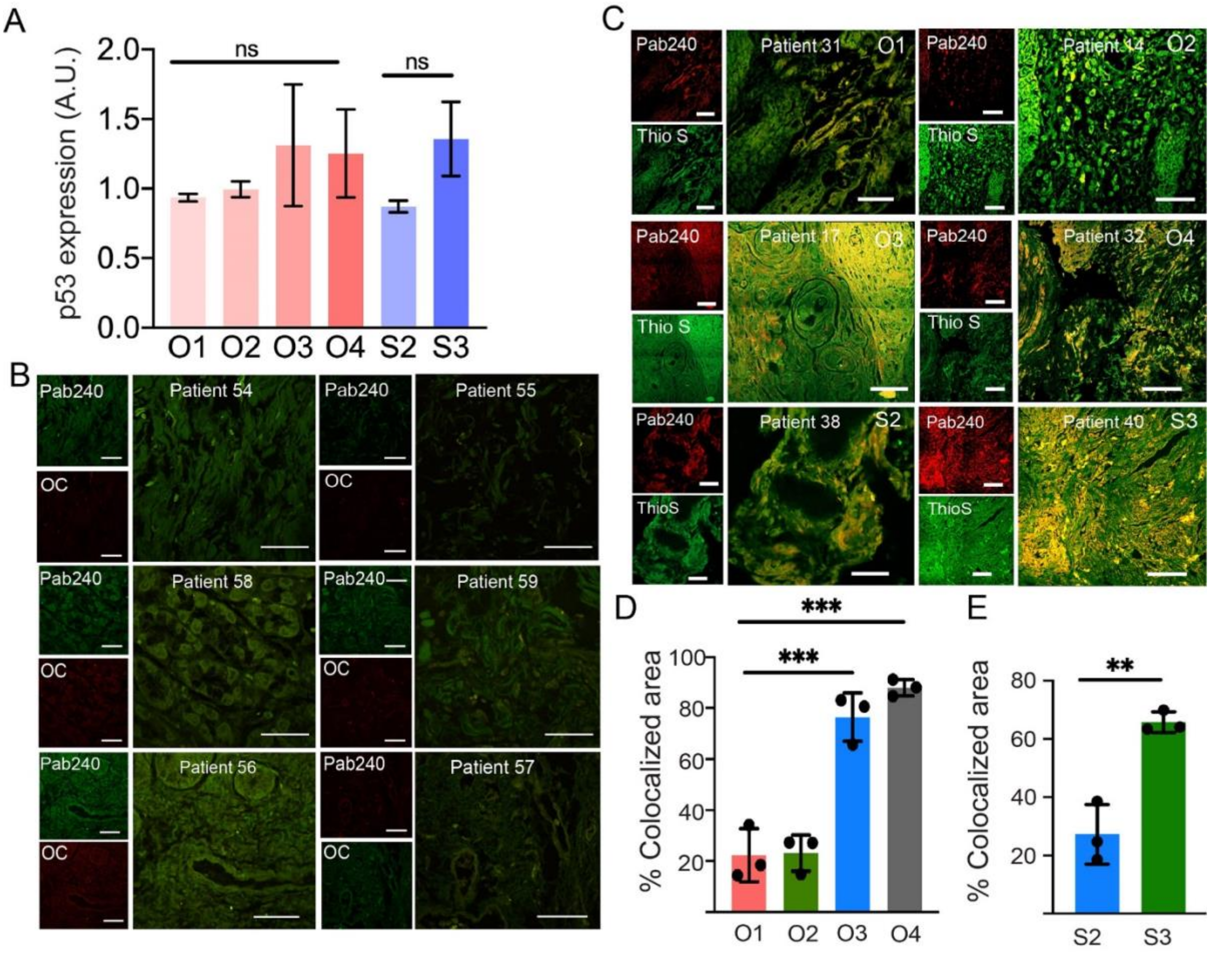


***Fig. S7*. Expression, misfolding and amyloid state of p53 in normal and cancer tissues.** a. Western blot quantification of p53 expression with respect to the GAPDH (loading control) showing similar expression in all the different grades of oral and stomach cancer tissues. The values were plotted as mean ± s.e.m., n=2 independent experiments. b. Double immunofluorescence study using Pab240 antibody (Santacruz, 1:500 dilution) and amyloid-specific antibody, OC (Abcam, 1:500 dilution) in oral and stomach normal tissues showing the absence of misfolded p53 and also co-localization of misfolded p53 and OC signals. Scale bars are 50 μm. c. Double immunofluorescence study of p53 using Pab240 antibody ((Santa Cruz Biotechnology, Dallas, TX, USA), 1:500 dilution) and amyloid specific dye, ThioS showing the co-localization of misfolded p53 and amyloid state (ThioS signals) in oral and stomach tissues. The data indicate the presence of p53 in amyloid state in human oral (O1, O2, O3 and O4) and stomach tumor tissues (S2 and S3). d,e. Co-localization analysis was performed using Image J analysis showing increase in the extent of co-localization between misfolded p53 with ThioS dye with higher grade of oral cancers (d) and stomach cancer tissues (e). Scale bars are 50 μm. The values were plotted as mean ± s.e.m., n=3 independent experiments.

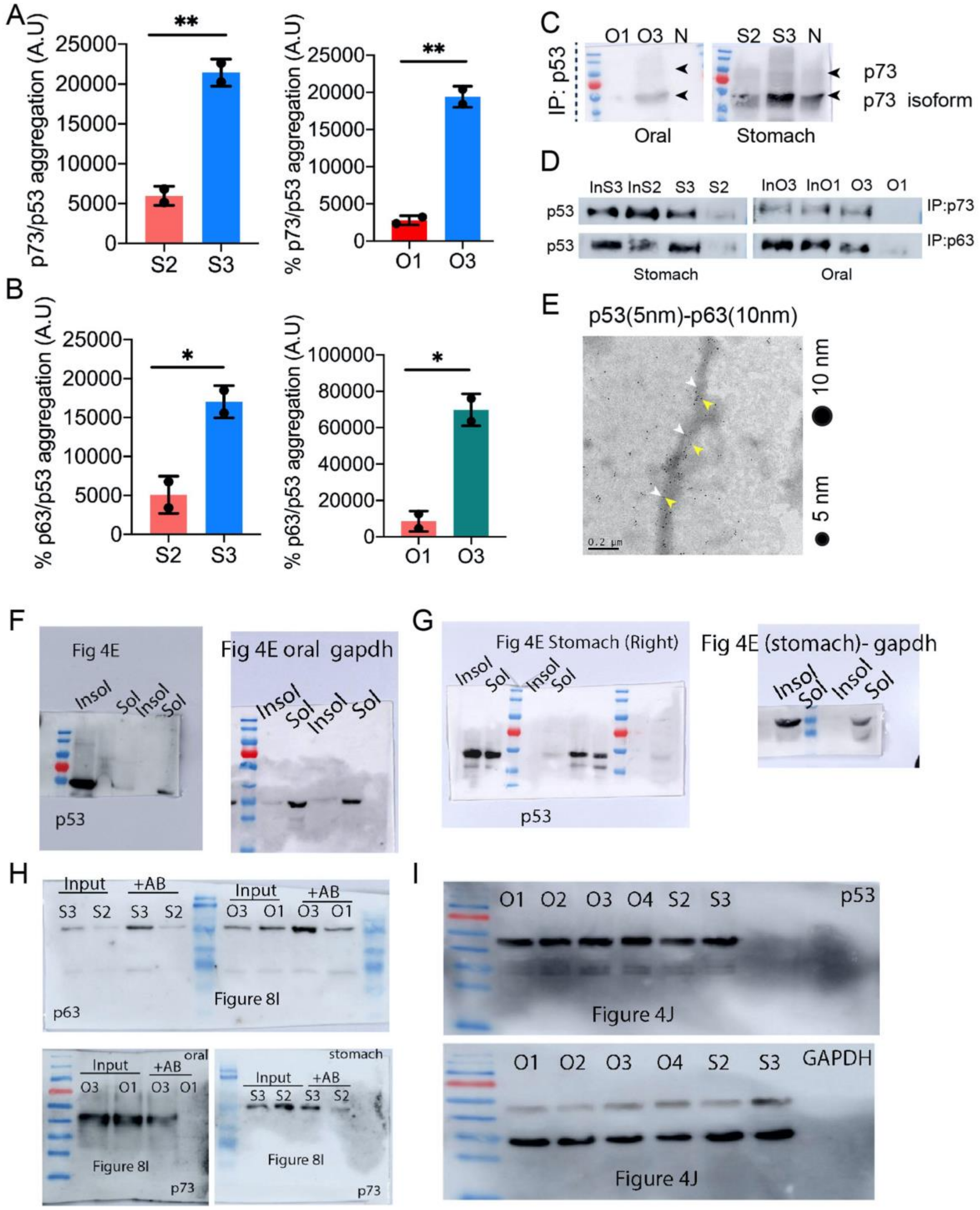


***Fig. S8. Quantification of the extent of p53 co-aggregation with p73 and p63 using western blot analysis.*** (a). Image J analysis of western blots determining the amount of p73, co-immunoprecipitated with p53 using anti-p53 Pab240 antibody from the various grade tumor tissue ((Oral Grade I (O1), III (O3) & Stomach Grade II (S2), III (S3)) tissue lysate. The data showing higher amount of p73 co-aggregated with p53 in the higher grade of stomach cancer (left panel) and oral cancer (right panel). In both the cancer types, p73 was observed to be co-aggregated to a significantly greater extent, with p53 in the higher cancer grades than in the lower grades. The values were plotted as mean ± s.e.m., n=3 independent experiments. (b). Quantification of p63 band intensity using Image J for both stomach (left panel) and oral (right panel) cancer tissues. In both the cancer types, p63 was observed to be co-aggregated to significantly greater extent with p53 in the higher cancer grades than in the lower grades.

The values were plotted as mean ± s.e.m., n=2 independent experiments. c. Additional Western blot analysis of immunoprecipitated p53 from the stomach and oral cancer tissues using p73 antibody showing the faint expression of full length p73 in immunoprecipitated p53. The low expression of p73 was observed in the higher cancer grade only. However, this was not observed in some of the patient. d. Western blot analysis of immunoprecipitated p73 and p63 from the stomach and oral cancer tissues showing higher expression of p53 when developed with p53 antibody in higher cancer grades. e. Uncropped Co-immunoelectron microscopy showing 5 nm and 10 nm gold particle decorations on the tissue isolated fibrils showing the presence of p53 and p63 in the fibrils isolated from oral tissue of a patient. Scale bars, 200 nm. F-I. Blot transparency image. All the original uncropped blots for the soluble and insoluble fraction of tissue lysate from oral an (F) stomach cancer (G) biopsies which is represented in the main manuscript Fig 4E. H. All original and uncropped blots for immunoprecipitated samples developed by p63 (above panel) and p73 (lower panel )antibodies from oral and stomach cancer tissues which is represented in the Fig. 8I of the main manuscript. I. Original and uncropped blot showing p53 expression across all the cancer grades developed by anti p53 antibody (above) and GAPDH (below) as a loading control which is represented in the main manuscript figure 4J.

***Table S1. p53 mutations detected by next-generation sequencing in oral and stomach cancer tissues.***

| P. No. | Patient ID | Tissue Type | Grade | Mutation type | Mutation | Clinical significance | Aggregated p53 localization |
|---|---|---|---|---|---|---|---|
| 1 | 500558 | Oral (Lower Lip) | II | Not Known | Not Known | NA | Nuclear |
| 2 | 500578 | Oral (RMT) | III | Not Known | Not Known | NA | Cytoplasmic |
| 3 | 500948 | Oral (Maxilla) | III | SNV | R175H | Pathogenic | Nuclear and cytoplasmic |
| | | | | SNV | P72R | Benign; Drug response mutation | |
| 4 | 500635 | Oral (FOM) | I | Not Known | Not Known | NA | Nuclear |
| 5 | 500903 | Oral (Alveolus | IV | Stop gained | R306/* | pathogenic | Nuclear |
| | | | | SNV | P72R | Drug response mutation | |
| 6 | 500536 | Oral (Lower Lip) | I | Stop gained | R196/* | Pathogenic | Nuclear and cytoplasmic |
| | | | | Deletion | PHHERC/- | NA | |
| | | | | SNV | P72R | Drug response Mutation | |
| 7 | 500275 | Oral (BM) | I | Not Known | Not Known | NA | Nuclear |
| 8 | 501027 | Oral (Alveolus) | IV | SNV | Intron | NA | Cytoplasm |
| | | | | Insertion | intron | NA | |
| | | | | SNV | E286V | Pathogenic | |
| | | | | Deletion | NA | NA | |
| | | | | SNV | P72R | Drug response mutation | |
| 9 | 500597 | Oral (BM) | III | Insertion | T102NX | NA | Nuclear |
| 10 | 501013 | Oral (Lower alveolus | IV | SNV | R267W | Likely pathogenic | Nuclear |
| | | | | SNV | R248W | Pathogenic | |
| | | | | SNV | P72R | Drug response mutation | |
| 11 | 501019 | Oral (Bucccal mucosa) | IV | SNV | R337C | Pathogenic | Nuclear |
| | | | | SNV | P72R | Drug response mutation | |
| 12 | 500886 | Oral (L. alveolus) | II | Stop gained | Q317/* | Unclear significance | Nuclear and cytoplasmic |
| | | | | SNV | P72R | Drug response mutation | |
| 13 | 500753 | Oral (BM) | I | SNV | P72R | Benign; Drug response mutation | Nuclear |
| 14 | 500618 | Oral (FOM) | II | Not Known | Not Known | NA | Nuclear |
| 15 | 500607 | Oral | II | Deletion | E/X | NA | Nuclear |
| | | | | Insertion | -/S | NA | |
| | | | | SNV | P72R | Drug response mutation | |
| 16 | 500600 | Oral (BM) | I | SNV | P72R | Drug response mutation | Nuclear and cytoplasmic |
| 17 | 500664 | Oral (BM) | III | Stop gained | R213/* | Pathogenic | Nuclear |

| | | | | | | | |
|---|---|---|---|---|---|---|---|
| | | | | SNV | P72R | Drug response mutation | |
| 18 | 500093 | Oral (Lower Lip) | II | SNV | P72R | Drug response mutation | Nuclear |
| 19 | 500083 | Oral (Upper Lip) | II | SNV | R282W | Loss and Gain of function | Nuclear and cytoplasmic |
| | | | | SNV | P72R | Benign; Drug response mutation | |
| 20 | 500783 | Oral (Upper Lip) | I | Not known | Not known | NA | Nuclear and Cytoplasm |
| 21 | 500740 | Oral (GBS) | II | Not known | Not known | NA | Nuclear and Cytoplasmic |
| 22 | 500190 | Oral (Upper alveolus) | II | Deletion | NA | NA | Cytoplasmic |
| | | | | SNV | P72R | Drug response mutation | |
| 23 | 500152 | Oral (Lower Lip) | II | Not known | Not known | NA | Nuclear and cytoplasm |
| 24 | 500352 | Oral | III | SNV | R282W | Pathogenic | Nuclear and Cytoplasmic |
| 25 | 500199 | Oral (Mandible) | III | Sop gained | R306/* | Pathogenic | Cytoplasmic |
| 26 | 500745 | Oral (Mandible) | II | Not known | Not known | NA | Nuclear and Cytoplasmic |
| 27 | 500066 | Oral (BM) | II | SNV | E285K | Temperature sensitive mutation; Loss of function | Nuclear and Cytoplasmic |
| | | | | SNV | P72R | Benign; Drug response mutation | |
| 28 | 500096 | Oral (Lower Lip) | II | SNV | P72R | Drug response mutation | Nuclear and Cytoplasmic |
| | | | | SNV | C176F | Pathogenic; Studied in HNSCC cells; LOF | |
| 29 | 500203 | Oral (L. alveolus) | II | SNV | P72R | Benign; Drug response mutation | Nuclear |
| | | | | Deletion | PSWPL/PX | Likely Benign | |
| 30 | 500530 | Oral (RMT) | II | Stop gained | R342/* | Pathogenic | Nuclear |
| | | | | SNV | P72R | Drug response mutation | |
| 31 | 500759 | Oral (L. alveolus) | I | SNV | R248L | Likely pathogenic | |
| | | | | SNV | P72R | Benign; Drug response mutation | |

| | | | | | | | |
|---|---|---|---|---|---|---|---|
| 32 | 501227 | Oral (Alveolus) | IV | SNV | INTRON | NA | Nuclear |
| | | | | SNV | R273H | Pathogenic | |
| | | | | SNV | P72R | Drug response mutation | Nuclear |
| 33 | 500124 | Oral (Lower Lip) | I | Intron SNV | NA | NA | |
| | | | | Deletion | RNSFE/X | NA | |
| | | | | Synomynous Variant | T | Benign | |
| | | | | SNV | P72R | Drug response mutation | |
| 34 | 500160 | Oral (GBS) | III | Stop gained | R342/* | Pathogenic | Nuclear |
| | | | | SNV | P72R | Drug response mutation | |
| 35 | 500201 | Oral (L. alveolus) | II | No Mutation | No Mutation | NA | Nuclear |
| 36 | 500118 | Oral (BM) | IV | SNV | P72R | Drug response mutation | Not Studied |
| 37 | 501352 | Stomach | II | SNV | P72R | Drug response mutation | Nuclear |
| 38 | 501432 | Stomach | II | No Mutation | No Mutation | NA | Cytoplasmic |
| 39 | 500326 | Stomach | III | SNV | G245S | Hotspot structural mutant | Nuclear and cytoplasmic |
| | | | | SNV | P72R | Drug response mutation | |
| 40 | 500874 | Stomach | III | Frame shift | E343X | NA | Nuclear and cytoplasmic |
| | | | | Insertion | -/S | NA | |
| | | | | SNV | R248W | Pathogenic | |
| | | | | SNV | P72R | Benign; Drug response mutation | |
| 41 | 501239 | Stomach | III | Not known | Not known | NA | Nuclear |
| 42 | 500636 | Stomach | III | No Mutation | No Mutation | NA | Cytoplasmic |
| 43 | 500065 | Stomach | II | SNV | P72R | Benign; Drug response mutation | Cytoplasmic |
| 44 | 601477 | Stomach | II | Deletion | NA | NA | Nuclear |
| | | | | SNV | P72R | Drug response mutation | |
| 45 | 500226 | Stomach | III | Intron SNV | NA | NA | Nuclear |
| | | | | SNV | I232F | Reported LOF | |
| | | | | SNV | P72R | Drug response mutation | |
| 46 | 500567 | Stomach | III | SNV | Y220H | Likely pathogenic; Structural analysis studied | Nuclear |
| | | | | SNV | A138S | Expressed in cell (50% activity) | |
| 47 | 601695 | Stomach | II | SNV | P72R | Benign; Drug response mutation | Nuclear and cytoplasmic |
| 48 | 601665 | Stomach | III | SNV | P72R | Benign; Drug response mutation | Nuclear and cytoplasmic |
| 49 | 501280 | Stomach | III | Synonynous variant | L | Benign | Nuclear and cytoplasmic |
| | | | | SNV | G245V | Pathogenic | |

| | | | | | | | |
|---|---|---|---|---|---|---|---|
| | | | | SNV | P72R | Benign; Drug response mutation | |
| 50 | 500450 | Stomach | III | SNV | C176F | Pathogenic; Studied in HNSCC cells; LOF | Cytoplasmic |
| | | | | SNV | P72R | Drug response mutation | |
| 51 | 601493 | Stomach | III | SNV | P72R | Benign; Drug response mutation | Cytoplasmic |
| 52 | 500067 | Stomach (GE Junction) | III | SNV | P72R | Benign; Drug response mutation | Nuclear |
| 53 | 500116 | Stomach | III | SNV | P72R | Drug response mutation | Nuclear |
| 54 | F | Oral | Normal | Not known | Not known | NA | NA |
| 55 | D | Oral | Normal | Not known | Not known | NA | NA |
| 56 | C14 | Oral | Normal | Not known | Not known | NA | NA |
| 57 | L | Oral | Normal | Not known | Not known | NA | NA |
| 58 | 1665 | Stomach | Normal | Not known | Not known | NA | NA |
| 59 | 567 | Stomach | Normal | Not known | Not known | NA | NA |

***Table S2. List of primers used in this study for ChIP assay.***

| | |
|---|---|
| p21 response element: | 5’ GTGGCTCTGATT GGCTTTCTG 3’ and 5’ CTGAAAACAGGCAGCCCAAG 3’ |
| PIG response element: | 5’ CAACGGCTCCTTTCTCTTCT 3’ and 5’ CCAGGCTTTTGGCACATTTA 3’ |
| GADD45 response element | 5’ TGTGGTACAGAACATGTCTAAGC 3’ and 5’ TGCAGATGTAGGTAGGGAGTAG 3’ |
| GSL2 response element | 5’ AGCCAAATAAGCCCTCCAACCC 3’ and 5’ TGTGGTTTCGCCATATCGGT 3’ |
| MDM2 response element | 5’ GTCAAGTTCAGACACGTTC 3’ and 5’ CCTCCAATCGCCACTGAACAC 3’ |

**References.**


1 Navalkar A, Pandey S, Singh N, Patel K, Datta D, Mohanty B *et al.* Direct evidence of cellular transformation by prion-like p53 amyloid infection. *J Cell Sci* 2021; **134**. doi:10.1242/jcs.258316.

2 Navalkar A, Paul A, Sakunthala A, Pandey S, Dey AK, Saha S *et al.* Oncogenic gain of function due to p53 amyloids occurs through aberrant alteration of cell cycle and proliferation. *J Cell Sci* 2022; **135**: jcs259500.
